# Uncharacterized protein c17orf80: a novel interactor of human mitochondrial nucleoids

**DOI:** 10.1101/2022.11.21.516320

**Authors:** Alisa Potter, Anu Hangas, Steffi Goffart, Martijn A. Huynen, Alfredo Cabrera-Orefice, Johannes N. Spelbrink

## Abstract

Molecular functions of many human proteins remain unstudied, despite the demonstrated association with diseases or pivotal molecular structures, such as mitochondrial DNA (mtDNA). This small genome is crucial for proper functioning of mitochondria, the energy-converting organelles. In mammals, mtDNA is arranged into macromolecular complexes called nucleoids that serve as functional stations for its maintenance and expression. Here, we aimed to explore an uncharacterized protein c17orf80, which was previously detected close to the nucleoid components by proximity-labelling mass spectrometry. To investigate the subcellular localization and function of c17orf80, we took an advantage of immunofluorescence microscopy, interaction proteomics and several biochemical assays. We demonstrate that c17orf80 is a mitochondrial membrane-associated protein that interacts with nucleoids even when mtDNA replication is inhibited. In addition, we show that c17orf80 is not essential for mtDNA maintenance and mitochondrial gene expression in cultured human cells. These results provide a basis for uncovering the molecular function of c17orf80 and the nature of its association with nucleoids, possibly leading to new insights about mtDNA and its expression.

## Introduction

Mitochondria are eukaryotic organelles responsible for aerobic metabolism and involved in regulation of the majority of cellular processes (Nunnari and Suomalainen, 2012). These organelles contain DNA molecules, i.e. mitochondrial DNA (mtDNA), that despite being small, are critical for energy conversion as they encode several subunits of oxidative phosphorylation (OXPHOS) complexes, as well as mitochondrial ribosomal and transfer RNAs. In a human mitochondrion, mtDNA exists in multiple circular copies organized in nucleic acid-protein complexes termed nucleoids, which individually resemble bacterial chromosomes (Lee and Han, 2017). The main structural protein component of nucleoids is the mitochondrial transcription factor A (TFAM), which fully coats mtDNA to tightly pack it into ellipsoid nanometric structures (Kukat et al., 2015). A variety of proteins associate with mtDNA to mediate its replication, repair, transcription, and distribution within the mitochondrial network. Such interactors involve, among others, core replication and transcription factors, namely DNA polymerase gamma (POLγ), mitochondrial single-strand binding protein (mtSSB), mitochondrial replicative helicase Twinkle, and mitochondrial RNA polymerase (POLRMT). While in the last decades the major components of nucleoids have been described, new proteins with functional roles in mtDNA-related processes are still regularly unveiled (Hensen et al., 2014).

In a previous study, we investigated the proximal interactome of Twinkle (Hensen et al., 2019), a core nucleoid protein. Along with the well-known nucleoid-associated proteins involved in mtDNA replication and transcription, we detected an uncharacterized protein, annotated as c17orf80, in close vicinity to Twinkle. Other groups have also identified this protein while investigating interactors of mtDNA-binding proteins via proximity-labelling proteomics (Antonicka et al., 2020; Hangas et al., 2022; Jiang et al., 2019). The close spatial proximity of this unknown protein to mtDNA replication and transcription apparatuses prompted us to investigate c17orf80 in more detail.

To date, there is no information regarding the molecular function(s) of c17orf80, and clinically significant data are very limited. However, some genetic association studies linked c17orf80 to autism spectrum disorder (Laird and Maertens, 2021; Mencer et al., 2021; Shaath, 2013) as well as pancreatic (Makler and Narayanan, 2017) and prostate cancers (Cheng et al., 2018). Many orphan proteins detected in genetic screens remain uncharacterized, as their molecular functions have not yet been investigated and are generally difficult to predict (Kustatscher et al., 2022).

Here, we used several strategies to further investigate and validate the interaction between c17orf80 and mitochondrial nucleoids. As an initial step in exploring the molecular function of c17orf80, we analysed the effect of its depletion on mtDNA maintenance and mitochondrial gene expression in cultured human cells.

## Results

### C17orf80 is a vertebrate protein with two transmembrane helices and a ZnF motif

The human c17orf80 gene (chromosome 17 open reading frame 80) is located on locus 17q25.1 and is predicted to produce three splice variants annotated in UniProt (Wang et al., 2021). According to Human Protein Atlas (Uhlén et al., 2015), c17orf80 mRNA is ubiquitously expressed in all human tissues, with the highest levels detected in the testes during spermatogenesis. The designated canonical c17orf80 isoform consists of 609 residues and has a theoretical molecular mass of 67 kDa. According to AlphaFold2 (Jumper et al., 2021) and Phyre2 (Kelley et al., 2015), most of the c17orf80 polypeptide lacks a well-defined secondary structure, except for the N- and C-termini **(Fig. 1A)**, suggesting it to be an intrinsically disordered protein (IDP) (van der Lee et al., 2014), as also predicted by the disorder prediction algorithm IUPRED (Dosztányi et al., 2005) **(Fig. 1B)**. Nevertheless, the remote homology detection server HHPred (Zimmermann et al., 2018) predicted that the N-terminus of c17orf80 is homologous to a zinc finger motif (ZnF, *E* = 0.1), and the C-terminus to subunit f of the mitochondrial F_1_F_O_-ATP synthase (ATP5MF, *E* = 2.6E-15) **(Fig. S1)**. The ZnF of c17orf80 has an arrangement of Cys-X2-Cys-X9-His-X3-Cys and belongs to the CCHC type typically involved in the binding of DNA or other proteins (Bijlmakers et al., 2016; Kim and Hudson, 1992). Because the ZnF spans only ∼20 amino acids, the *E* value is relatively high; however, the cysteine and histidine residues as well as some other residues of that region are well conserved among the c17orf80 orthologs **(Fig. 1C and 1D)**. The homology with the ATP synthase subunit f covers almost that complete protein, including its two transmembrane (TM) helices that correspond to the helices predicted in c17orf80 **(Fig. S1)**. Note that both the N- and C-termini of the ATP synthase subunit f are located in the mitochondrial matrix (Gu et al., 2019). If this topology is preserved in c17orf80, this suggests that it associates with the inner mitochondrial membrane with most of its polypeptide in the matrix.

**Figure 1.**
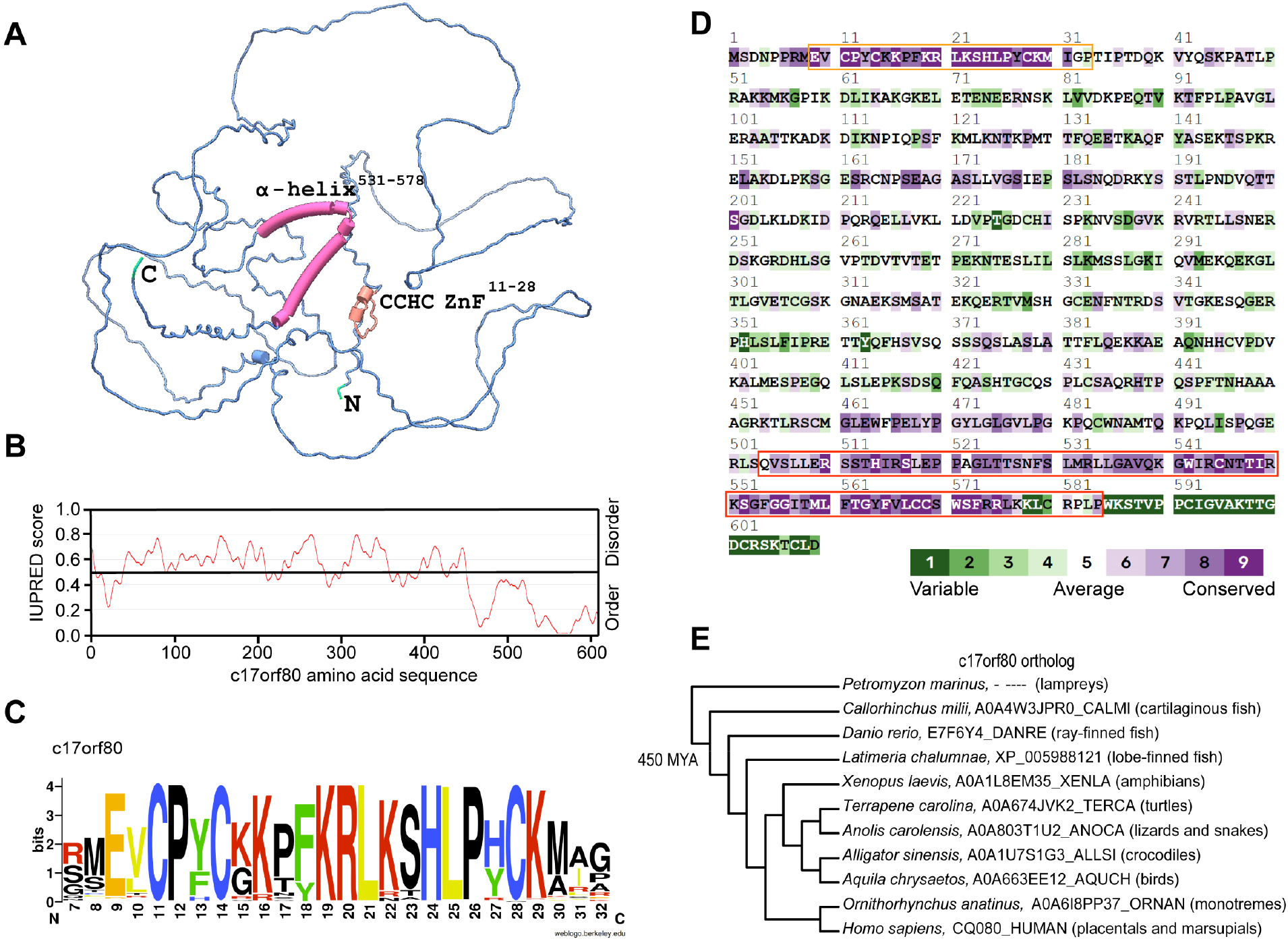
C17orf80 lacks a defined secondary structure but contains a ZnF motif and a two TM helices. **A**. AlphaFold2 predicted structure of c17orf80. Most of the protein is predicted to be unstructured, except for the N- and C-termini. **B**. Approximately 80% of c17orf80 protein is intrinsically disordered as predicted by IUPRED with a cutoff for disorder of 0.5. **C**. Sequence logo of the N-terminus of c17orf80 shows conservation of the cysteine (C) and histidine (H) amino acids among others. **D**. Evolutionary conservation analysis. The N- and C-termini of c17orf80 are the most conserved and homologous to a zinc finger (orange box) and the subunit f of the F_1_F_O_-ATP synthase (red box). **E**. Phylogenetic distribution of c17orf80, as deduced from a JackHMMER search against UniProt. Protein IDs from representative species of the various vertebrate clades are indicated. C17orf80 orthologs, which are defined by having both an N-terminal ZnF and a C-terminal domain homologous to F_1_F_O_-ATP synthase subunit f, cannot be detected outside the gnathostome vertebrates that arose about 450 million years ago (MYA).

To determine the evolutionary origin of the c17orf80 orthologous group, we used JackHMMER (Johnson et al., 2010) to detect homologs that contain both the N-terminal zinc finger and the C-terminal domain that is homologous to the ATP synthase subunit f. The most distantly related to humans species in which we detected full-length homologs of c17orf80 are cartilaginous fish, such as sharks (e.g. AOA4W3JPRO_CALMI in the Ghost shark). This makes c17orf80, relative to other mitochondrial proteins, a young gene that arose about 450 million years ago, early in the evolution of vertebrates after branching off from jawless vertebrates like Lampreys **(Fig. 1E)**.

### C17orf80 locates in mitochondria and associates with the inner membrane

C17orf80 was identified in BioID screens as a potential interactor of mitochondrial nucleoid-associated proteins. According to the DeepMito (Savojardo et al., 2019) and Mitofates (Fukasawa et al., 2015) prediction algorithms, c17orf80 has a low prediction score of localizing to mitochondria or for containing an N-terminal mitochondrial pre-sequence (0.17 and 0.045 for DeepMito and Mitofates, respectively). However, c17orf80 may contain an internal matrix-targeting sequence, as predicted by the iMTS-L algorithm (Schneider et al., 2021), with a propensity score of 3.91. C17orf80 is not annotated in MitoCarta 3.0 (Rath et al., 2021), a list of proteins with strong evidence of mitochondrial localization. This protein is nonetheless classified as “likely associated with mitochondria” in the Integrated Mitochondrial Protein Index (IMPI) collection (Smith and Robinson, 2018). The latter is consistent with several proteomics studies of subcellular fractions, showing that c17orf80 localizes in mitochondria (Antonicka et al., 2020; Morgenstern et al., 2021).

To confirm that c17orf80 locates in mitochondria, we used immunofluorescence (IF) microscopy. We detected co-immunofluorescence of c17orf80, mtDNA, and a mitochondrial marker cyclophilin D (CypD) or a nucleoid marker TFAM in human osteosarcoma cells (U2OS) **(Fig. 2A)**. We observed that the majority of the c17orf80 IF signal overlapped with the mitochondrial network, showing both uniform and punctate patterns. We quantified the colocalization using Manders coefficients (Manders et al., 1993) **(Fig. 2B)**. The Manders coefficient M1 calculated for c17orf80:CypD indicated that most of the c17orf80 IF signal colocalized with the mitochondrial network, while Manders coefficients M1 calculated for TFAM:c17orf80 and mtDNA:c17orf80 indicated that most of the nucleoids colocalized with c17orf80. In turn, Manders coefficients M2 for c17orf80:CypD, TFAM:c17orf80 and mtDNA:c17orf80 pairs indicated that only half of the c17orf80 signal colocalized with mitochondria or nucleoids. The M1 and M2 coefficients for mtDNA:CypD and mtDNA:TFAM pairs demonstrated the expected localization of mtDNA in the mitochondrial network and the exact colocalization of mtDNA and TFAM. These data confirmed that the majority of the cellular c17orf80 locates within the mitochondrial network.

**Figure 2.**
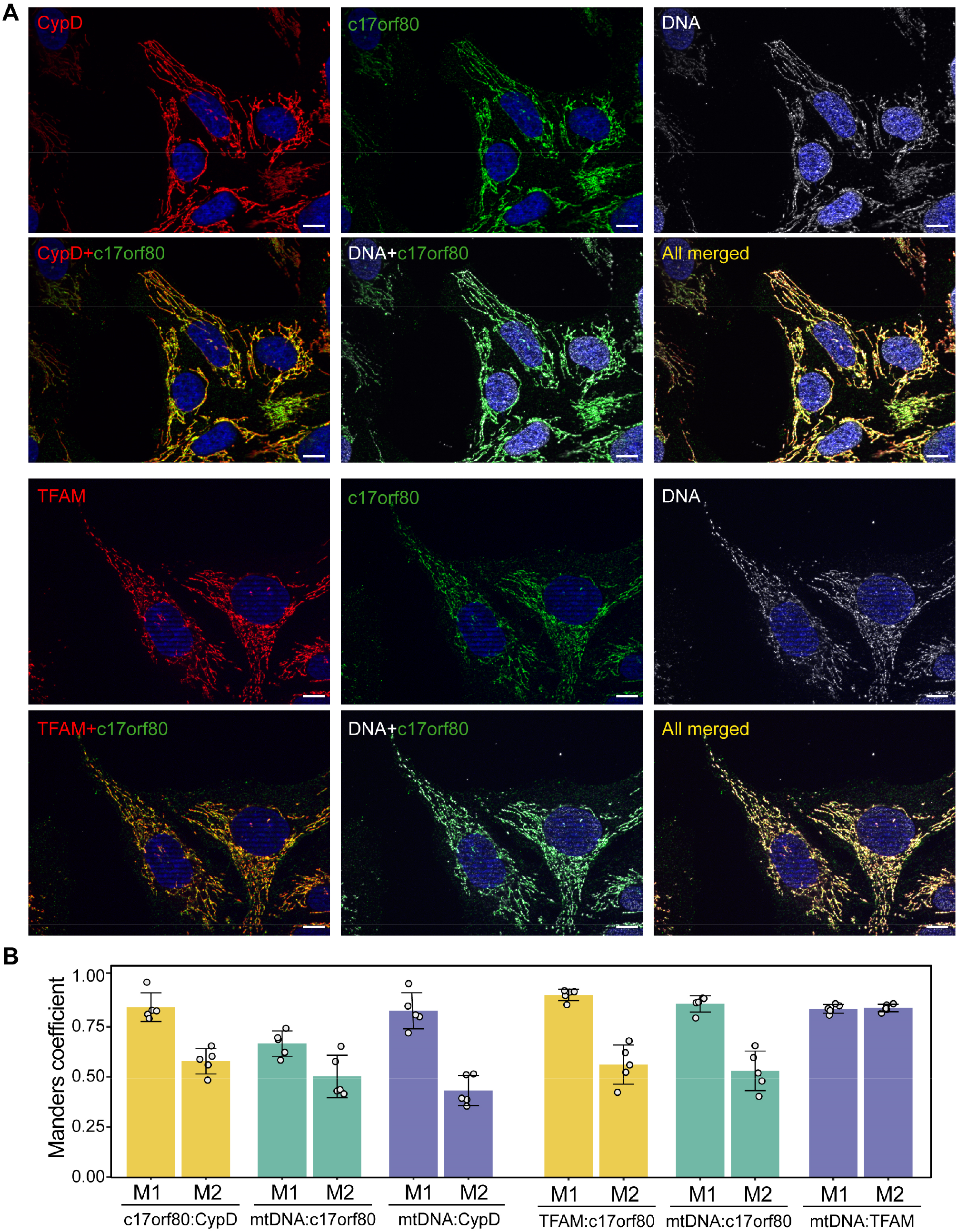
C17orf80 colocalizes with the mitochondrial network. **A**. Representative IF images of U2OS cells co-labelled with c17orf80, DNA, and CypD or TFAM antibodies. **B**. Quantification of Manders coefficients M1 and M2 for the indicated pairs of antibodies. Data are mean ± SD from five images with 2-6 cells per image with individual values overlaid. The area of nuclei was excluded from the analysis. The scale bar is 10 μm.

Additional IF signal obtained with the c17orf80 antibody was observed in the cytoplasm. To check whether this signal relates to c17orf80, we validated the specificity of the antibody using siRNA-mediated depletion and found that only the mitochondria-derived signal was specific to c17orf80 **(Fig. S2)**.

To further define the sub-mitochondrial localization of c17orf80, we treated fixed U2OS cells with either digitonin or a combination of digitonin and Triton X-100. The cells were then probed with antibodies against an outer mitochondrial membrane (OMM) protein TOM20, a matrix protein MRPL12, as well as DNA and c17orf80 **(Fig. 3A)**. The added amount of digitonin permeabilizes the plasma membrane and OMM while keeping the inner mitochondrial membrane (IMM) mostly intact. Consequently, this condition enables TOM20 to be detected by IF, but not MRPL12 or mtDNA. The addition of Triton X-100 disrupts both the OMM and IMM, thus making the IMM and matrix accessible for antibodies. Under these conditions, TOM20, MRPL12 and mtDNA can be detected. In this assay, the IF signal of c17orf80 was only detected in the cells treated with both detergents. Considering that the epitope for the c17orf80 antibody is located at the N-terminal part of the protein (35-261 aa), this result suggests that at least this region of c17orf80 resides in the internal mitochondrial compartments.

**Figure 3.**
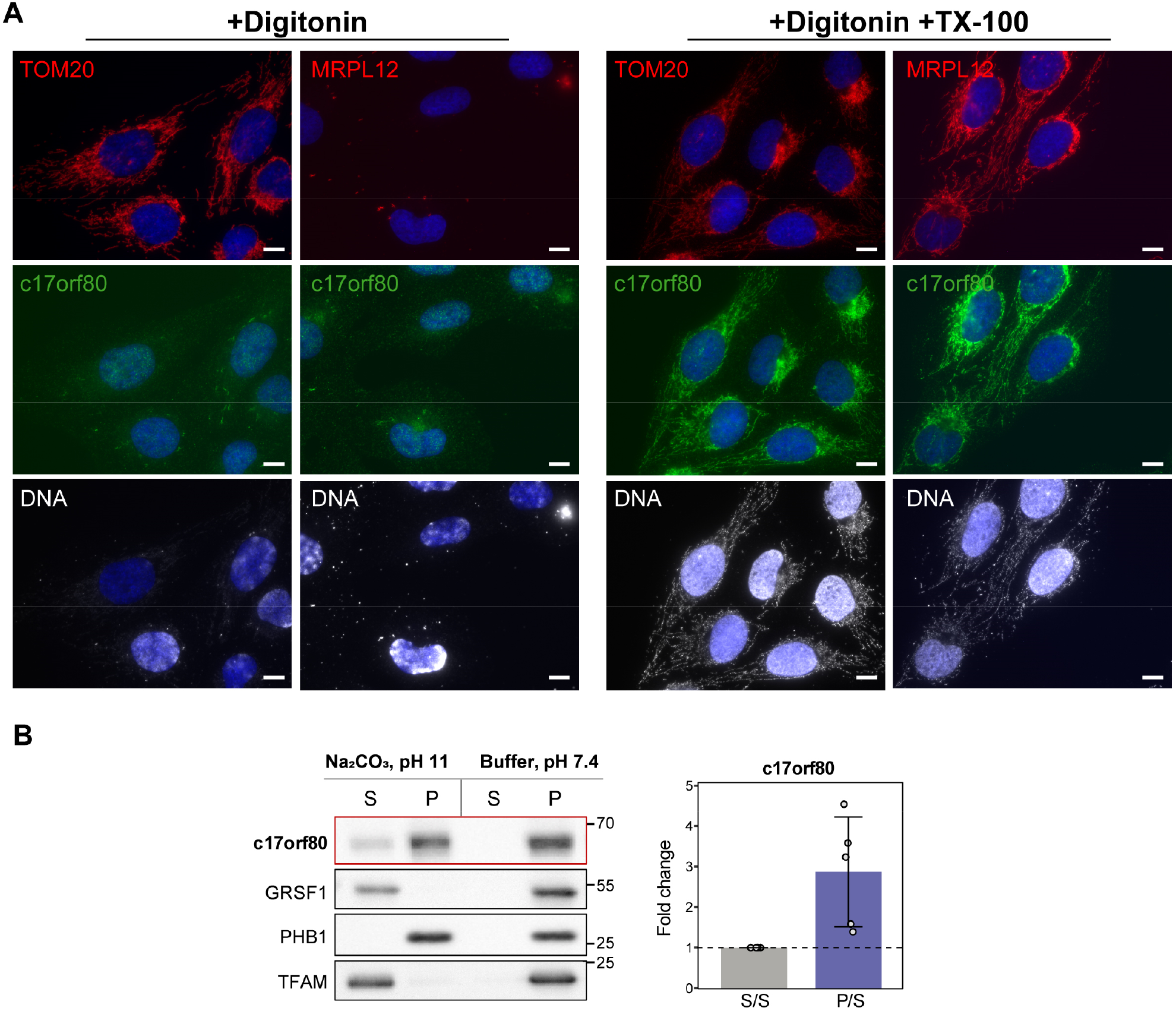
C17orf80 locates inside of mitochondria and associates with mitochondrial membranes. **A**. C17orf80 is inaccessible for antibody binding without permeabilization of mitochondrial membranes. IF images of U2OS cells permeabilized with either digitonin or a combination of digitonin and TX-100 and co-labelled with antibodies against c17orf80, DNA and TOM20 or MRPL12. The scale bar is 10 μm. **B**. C17orf80 pellets with the membrane fraction in sodium carbonate extraction. Western blots showing levels of c17orf80, PHB1 (integral membrane protein), GRSF1 and TFAM (soluble proteins) in pellet (P) and supernatant (S) fractions. The bar chart shows mean ± SD of the S-to-S and P-to-S ratios for c17orf80 from five independent experiments with overlaid individual values.

Since c17orf80 was predicted to contain C-terminal TM helices, we sought to determine whether the protein is membrane-associated. For this, we subjected intact mitochondria isolated from HEK293 cells to sodium carbonate alkaline extraction (Fujiki et al., 1982a; Fujiki et al., 1982b). This method allows the separation of integral membrane proteins from peripheral and soluble ones. In this experiment, the majority of c17orf80 was found in the pellet fraction. However, a visible portion of the protein also appeared in the soluble fraction and the distribution of c17orf80 between pellet and supernatant varied between the experiments **(Fig. 3B)**. This result agrees with the structural prediction of c17orf80 containing TM helices, nevertheless indicating that it is not an integral membrane protein.

Taken together, the results of the colocalization analysis and IF antibody accessibility test combined with alkaline extraction indicated that c17orf80 is a mitochondrial protein that is located in the mitochondrial matrix and associates with the inner mitochondrial membrane.

### C17orf80 colocalizes with mtDNA

As the previous c17orf80 BioID interaction data suggested a link between c17orf80 and mtDNA or mitochondrial gene expression (Hensen et al., 2019), we sought to determine whether c17orf80 associates with mitochondrial nucleoids or RNA granules, the RNA-processing structures that are often found adjacent to nucleoids (Antonicka et al., 2013; Jourdain et al., 2013). For this, we performed co-immunofluorescence experiments labelling nucleoids with antibodies against TFAM and/or DNA, and RNA granules with antibodies against 5-bromouridine (BrU) following BrU labelling. Because of variability in intensity and background of c17orf80 IF, we opted for qualitative colocalization studies using chemical treatments that affect mtDNA quantity and nucleoid morphology.

First, we investigated whether c17orf80 colocalizes with mitochondrial nucleoids or RNA granules under regular culturing conditions. To this end, we co-labelled U2OS cells with c17orf80, TFAM and DNA antibodies. We observed that c17orf80 formed foci that overlapped with mtDNA and TFAM **(Fig. 4A)**. The RNA granules that appeared after 20 min of BrU labelling partially overlapped with nucleoids or were located next to them; expectedly, not all of the mtDNA spots had an adjacent RNA granule. We observed that c17orf80 preferentially colocalized with nucleoids but not with RNA granules **(Fig. 4B)**.

**Figure 4.**
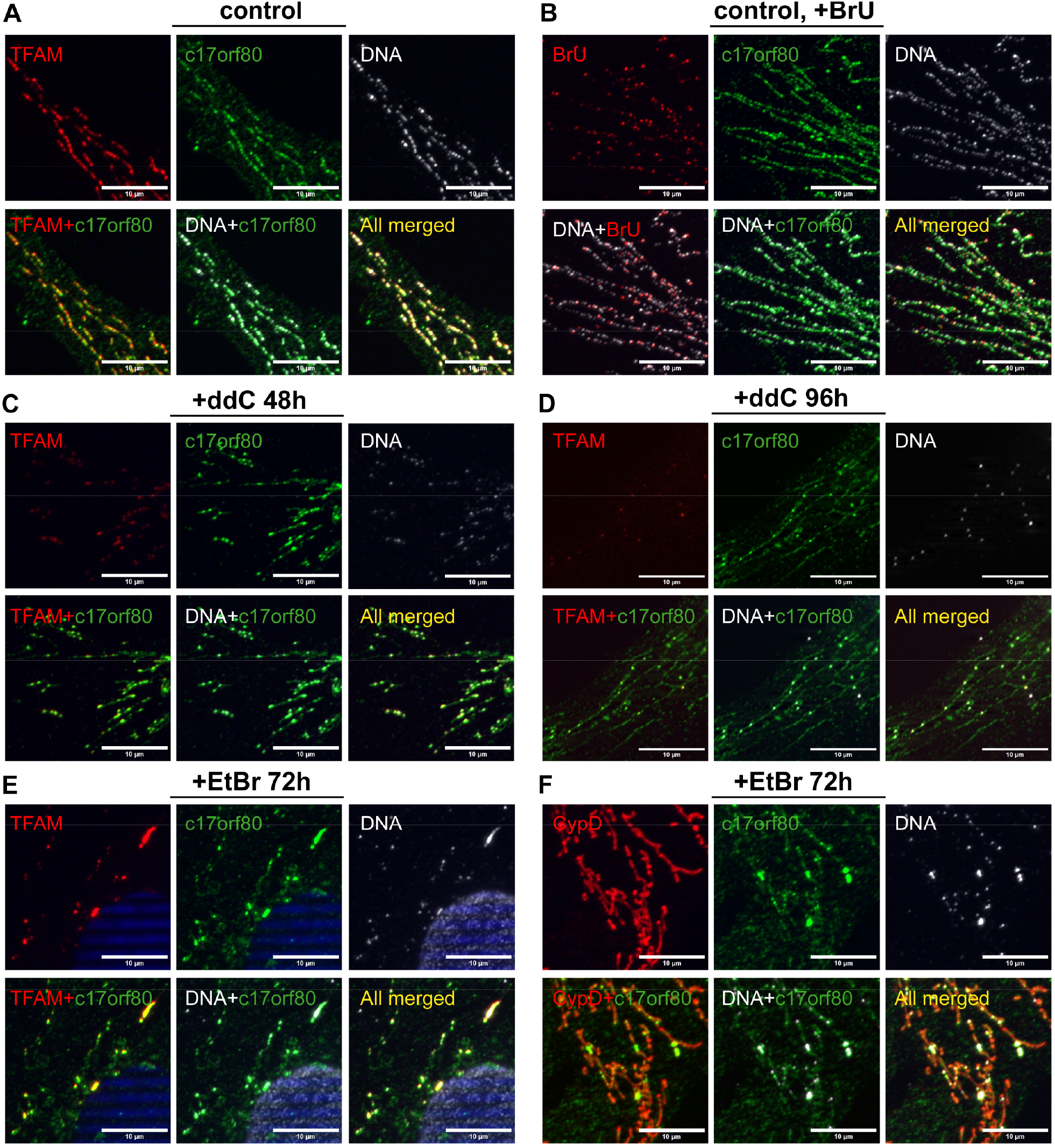
C17orf80 colocalizes with mtDNA. Co-immunostaining with c17orf80, TFAM (nucleoids), CypD (mitochondrial network), DNA (nucleoids and nuclei), or BrU (RNA-granules) antibodies. Nuclei stained with DAPI (blue). Zoomed-in sections of 25×25 μm are shown. The scale bar is 10 μm. For uncropped images, see **Suppl. Fig. S3. A**. The punctate signal of c17orf80 colocalizes with nucleoids in untreated cells. **B**. C17orf80 colocalizes with nucleoids and only partially with RNA granules. Cells were treated with 2.5 mM BrU for 20 min. **C**. After partial mtDNA depletion with ddC, c17orf80 still colocalizes with nucleoids. Cells were treated with 100 μM ddC for 48 h. **D**. After almost complete depletion of mtDNA, most of the c17orf80 signal is diffused, but a portion of it still colocalizes with the remaining nucleoids. Cells were treated with 100 μM ddC for 96 h. **E-F**. C17orf80 colocalizes with the enlarged nucleoids that appear after EtBr treatment (E). Clustered nucleoids do not represent individual fragmented mitochondria in EtBr-treated cells (F). Cells were treated with 50 ng/ml EtBr for 72 h.

To examine whether c17orf80 foci were dependent on presence of mtDNA, we depleted mtDNA by treating cells with 2’-3’-dideoxycytidine (ddC) for two or four days. After depletion, the cells were co-labelled with c17orf80, TFAM, and DNA antibodies. After two days of ddC treatment, partial depletion of mtDNA was observed, while c17orf80 was still detected in foci overlapping with nucleoids. Moreover, the c17orf80 IF signal visibly increased, whereas the signal of TFAM was reduced **(Fig. 4C)**. After four days of ddC treatment, c17orf80 was still found in foci overlapping with the remaining nucleoids, although most of its signal was uniformly distributed along the mitochondrial network **(Fig. 4D)**.

As we observed c17orf80 to remain in nucleoids after mtDNA depletion with ddC, we tested whether the same was true when depleting mtDNA with ethidium bromide (EtBr). While ddC primarily affects mitochondrial gene expression by blocking mtDNA elongation thus resulting in mtDNA-loss (Nelson et al., 1997), EtBr intercalates into mtDNA and actively hampers both replication and transcription (Coppey-Moisan et al., 1996; Hayakawa et al., 1998). We treated cells with a moderate concentration of EtBr for three days, which induced not only mtDNA depletion, but also frequent clustering of nucleoids, a phenomenon described previously (Alán et al., 2016). Accumulation of c17orf80 was observed in the clustered nucleoids along with TFAM. The remaining nucleoids, with normal morphology, also contained c17orf80 **(Fig. 4E)**. In parallel, we also detected the mitochondrial network with CypD to confirm that the clustered nucleoids are not just individual fragmented mitochondria **(Fig. 4F)**.

These data indicate that c17orf80 not only interacts with the nucleoids under regular conditions, but also remains associated after treatment with mtDNA replication inhibitors ddC and EtBr.

### C17orf80 accumulates in nucleoids upon a short treatment with ddC

In our ddC treatment experiment, we observed that c17orf80 foci appeared brighter after two days of the treatment. At the same time, IF signals of TFAM and DNA were expectedly lower in ddC-treated cells whereas nucleoid morphology seemed unaffected **(Fig. 5A)**. We calculated an average particle size and intensity levels (represented by object mean grey intensity) of the c17orf80, TFAM, and mtDNA spots using the ImageJ software. We found that while the size of TFAM and mtDNA spots decreased by 1.5- and 3-fold, respectively, after ddC treatment, the size of c17orf80 spots increased by 35% **(Fig. 5B)**. The calculated intensity change was very small for all three channels. However, the intensity of c17orf80 was significantly higher (by 11%) after ddC treatment, in contrast to a slight decrease for mtDNA and TFAM. Additional IF images demonstrating changes in c17orf80 IF upon ddC treatment are provided in high-resolution as supplementary data **(Suppl. Data 1)**.

**Figure 5.**
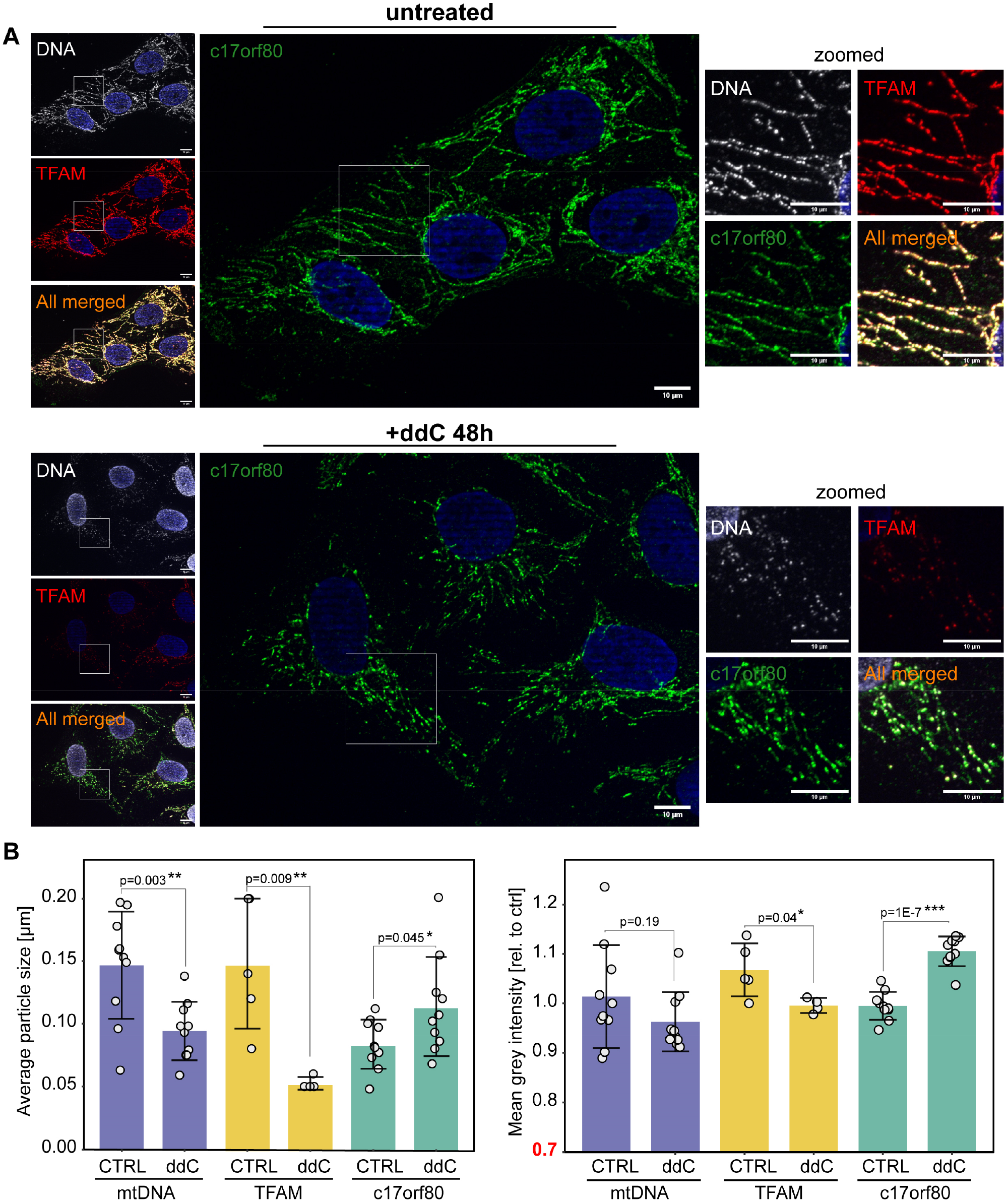
C17orf80 concentrates in nucleoids following treatment with ddC. **A**. Representative images of U2OS cells treated or not with 100 μM ddC for 48 h. Co-immunostaining with c17orf80, TFAM (nucleoids), and DNA (nucleoids and nuclei) antibodies. Nuclei stained with DAPI (blue). The scale bar is 10 μm. Signals of TFAM and mtDNA in ddC-treated cells were reduced, whereas the nucleoid-specific signal of c17orf80 was increased. **B**. Quantification of average particle size and grey intensity in control and ddC-treated cells. Data are shown as mean ± SD from ten mtDNA and c17orf80 images from three independent experiments, and four or five TFAM images from two independent experiments; individual values overlaid; unpaired Student’s t-test. Cutoffs for statistical significance: \**p* ≤ 0.05, \*\**p* ≤ 0.01, \*\*\**p* ≤ 0.001. The area of nuclei was excluded from the analysis.

To further examine if changes in c17orf80 localization could result from stress situations or mitochondrial translation blockage, cells were exposed to either H_2_O_2_ or UV light, and incubated with chloramphenicol, respectively (data not shown). We did not detect a similar change in c17orf80 IF signals as aforementioned, which suggests that its accumulation in nucleoids upon ddC treatment is a response to mtDNA replication/depletion stress and not to oxidative stress, mtDNA damage or translation inhibition.

As we thought that the increase in c17orf80 IF after ddC treatment could reflect its molecular function, we decided to explore whether any other nucleoid components behave similarly. Thus, we detected IF of TFAM, mtSSB, Twinkle, and POLRMT in control and ddC-treated cells; in addition, we tested the effect of ddC treatment on RNA granules **(Fig. S4)**. We observed that upon ddC treatment: a) the TFAM signal was reduced; b) the signal of mtSSB, a marker of active mtDNA replication (Rajala et al., 2014), was reduced and its punctate pattern became dispersed; c) Twinkle remained unchanged; d) the POLRMT signal was reduced and generally uniform even in control cells. The RNA granule formation was inhibited after two days of ddC treatment, but some uniform BrU staining was still observed, suggesting that mitochondrial transcription is not fully suppressed. It should be noted that we tested only proteins for which we had IF-compatible antibodies. Nevertheless, these results indicate that a short treatment with ddC does not induce a sudden enrichment of mitochondrial transcription and replication factors in nucleoids. Thus, the nature of c17orf80 concentration in the remaining nucleoids remains elusive, but our findings suggest a high affinity of this protein for mtDNA or nucleoid protein components.

### C17orf80 interacts with mitochondrial replication and gene expression factors

In a study of the mitochondrial protein interactome performed by Antonicka et al. (Antonicka et al., 2020), which employed a proximity-dependent biotinylation assay (BioID), c17orf80 was identified as prey for 59 out of 100 baits used, with the highest fold enrichment for TFAM, AUH, MTERFD1 and mtSSB baits. The authors also used c17orf80 as bait but the resulting hit list did not contain the above-mentioned proteins. In contrast, it mostly consisted of membrane-bound and membrane-associated proteins.

Here, we set out to investigate the protein interactors of c17orf80 in more detail, and performed a BioID assay using the Flp-In™ T-Rex™ cell system, allowing for stable and inducible expression of c17orf80 fused with BirA* (Roux et al., 2012) added to either the N- or C-terminus. Prior to the experiment, we confirmed that both N- and C-terminally tagged fusions of c17orf80-BirA* were localized to mitochondria. To this end, we used transient transfection of the fusion proteins in U2OS cells followed by IF detection **(Fig. S6)**. As a negative control for BioID pulldowns, we used the same cells cultured without addition of the inducing agent (anhydrotetracycline, AnTET).

We performed BioID pulldowns in quadruplicate using concentrations of AnTET optimized for each cell line individually. Cells were induced or not with AnTET and simultaneously supplemented with biotin to allow for biotinylation of all preys that c17orf80-BirA* fusions interact with after being synthesized in the cytoplasm. Biotinylated proteins were pulled down from total cell lysates with streptavidin-coated agarose beads, trypsin-digested on the beads, and analysed by mass spectrometry.

We first analysed which proteins were biotinylated with the N-terminally tagged c17orf80-BirA* fusion by comparing it to the pulldown performed with non-induced cells. The statistically significant differences between protein abundances were determined in the Perseus software (Tyanova et al., 2016) using a two-tailed Welch’s t-test with a permutation-based false-discovery rate (FDR) cutoff of 0.05 and artificial within-group variance (S0) set to 0.1 (Tusher et al., 2001). This analysis revealed 78 potential protein interactors of c17orf80 (hits), of which 40 were annotated in MitoCarta 3.0. At least 15 hits were involved in mtDNA replication or gene expression, with the highest enrichment for the core nucleoid proteins TFAM (12-fold), Twinkle (7-fold), and mtSSB (7-fold) **(Fig. 6A, 6C, Suppl. Data 2)**. This finding is in line with the previous proximity-labelling experiments that suggested that c17orf80 is located in close vicinity to mtDNA replication factors, as well as with our immunofluorescence data showing that c17orf80 colocalizes with mitochondrial nucleoids.

**Figure 6.**
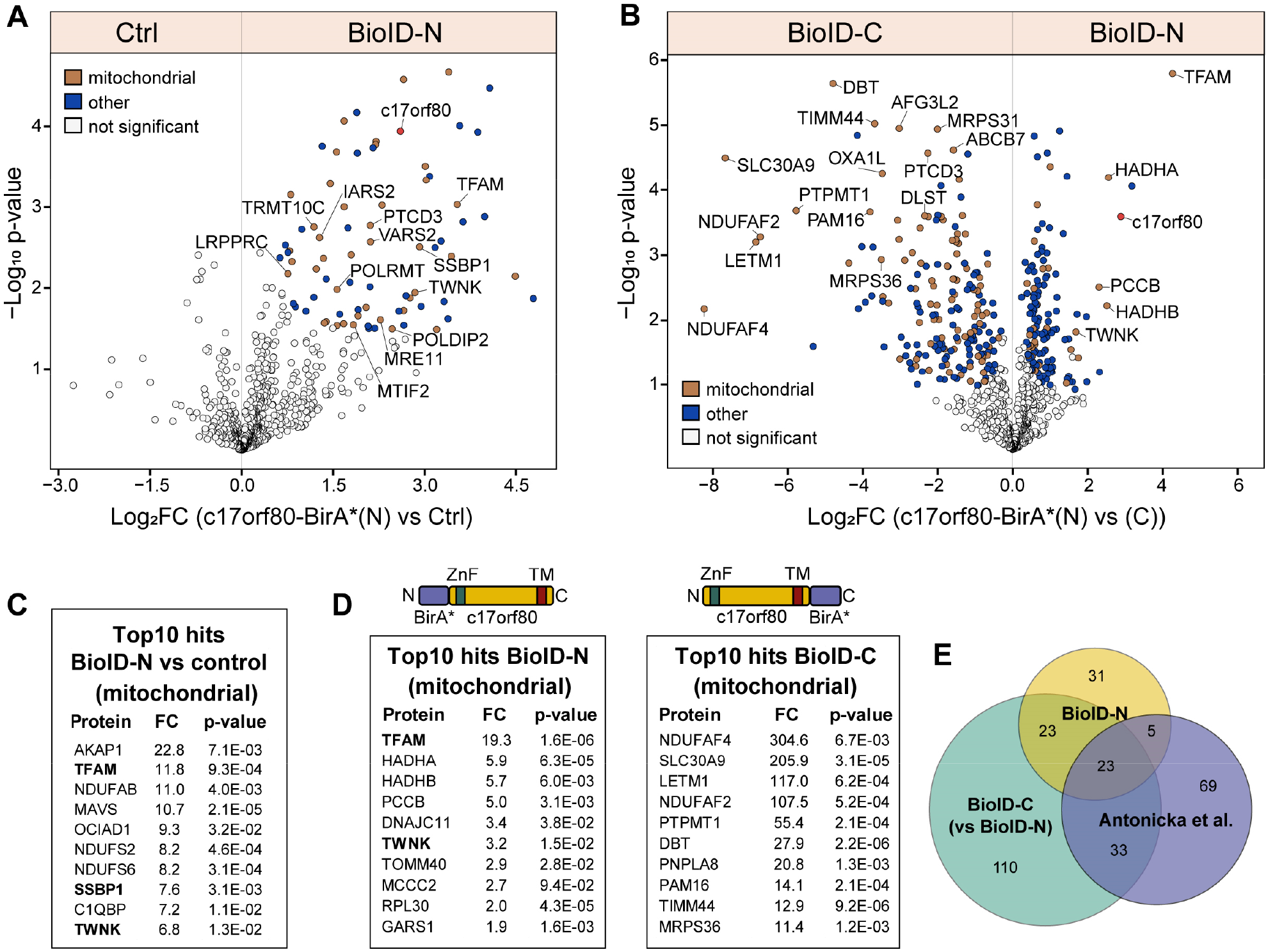
c17orf80 interacts with TFAM and other proteins involved in mitochondrial gene expression. A-B. Volcano plots depict average log_2_-transformed fold changes of protein abundances plotted against negative log_10_-transformed *p* values for four biological replicates. Statistical significance was determined by FDR < 0.05 with S0 = 0.1. **A**. N-terminal interactome of c17orf80. Fold change was defined as a ratio of c17orf80-BirA*(N) to control. Coloured points indicate proteins that were significantly enriched in the c17orf80-BirA*(N) pulldown. **B**. Comparison of proteins labelled by either N- or C-terminally tagged c17orf80-BirA* fusions. Fold change was defined as a ratio of c17orf80-BirA*(N) to c17orf80-BirA*(C). Coloured points indicate proteins that were significantly enriched in either of the pulldowns. **C-D**. Top10 hits annotated in MitoCarta 3.0 for each pulldown with the indicated fold change (FC) and *p* values. Above the tables of panel D: schematic representations of c17orf80-BirA* fusion proteins. **E**. Venn diagram representing the number of hits shared between the datasets presented here and the results of c17orf80 BioID obtained previously (Antonicka et al., 2020) (filtered proteins with BFDR = 0).

When we applied the same statistical analysis to the BioID pulldown performed with C-terminal c17orf80 BioID fusion, we noticed only a few preys reached the statistical significance threshold due to low fold-change values **(Fig. S6B)**. We determined that the non-induced control contained an equally large set of proteins as the BioID pulldown. In addition, this control had a higher overall protein abundance than the control for c17orf80-BirA*(N) **(Suppl. Data 1)**. It is possible that the amount of c17orf80-BirA*(C) fusion protein produced by leaky expression in non-induced cells was sufficient to biotinylate as many preys as in the induced ones, even though the protein level was 9-fold higher upon induction **(Fig. S6A)**.

Nevertheless, a direct comparison of BioID pulldowns performed with C- and N-terminally tagged c17orf80-BirA* fusions showed distinct protein identifications. The C-term BioID pulldown was more enriched for mitochondrial inner membrane proteins, such as subunits of the TIM complex, Complex I, BCKDC, as well as OXAL1 and LETM1 **(Fig. 6B, 6D)**. This hit list was similar to that published previously for c17orf80 C-terminal BioID (Antonicka et al., 2020), with 56 shared hits **(Fig. 6E)**. In contrast, in the N-term BioID pulldown, we identified a 19-fold enrichment for TFAM and 3-fold enrichment for Twinkle compared to the C-term pulldown.

It should be noted that although the c17orf80-BirA*(C) fusion protein was imported into mitochondria, it, in contrast to c17orf80-BirA*(N), did not biotinylate the endogenous c17orf80, as evidenced by the absence of biotinylated c17orf80 after induction **(Fig. S6A)**. This might indicate that the C-terminal region of c17orf80 is actually located in the inner mitochondrial membrane, and when BirA* is added to the C-terminus, it cannot reach matrix proteins. Nonetheless, there is a possibility that the c17orf80-BirA*(C) fusion protein does not preserve the exact location and function of the endogenous c17orf80. Thus, the results of proximity-labelling with the C-terminally tagged c17orf80-BirA* fusion should be interpreted with caution.

### Complexome profiling identifies c17orf80 migrating at high molecular mass

Thus far we have shown that c17orf80 colocalizes with mitochondrial nucleoids and is located in close proximity to multiple nucleoid-interacting proteins. To further study these interactions and gain more evidence regarding the potential occurrence of c17orf80 in DNA-/RNA-associated protein complexes, we performed complexome profiling (CP) (Cabrera-Orefice et al., 2022; Van Strien et al., 2019; Wessels et al., 2013). CP is an unbiased approach that combines the separation of native proteins and protein complexes, typically by gel electrophoresis, with quantitative tandem MS identification of individual fractions followed by data clustering. Individual signals from components of protein complexes typically co-fractionate, hence showing similar intensity patterns across gel fractions. In contrast to BioID, CP does not require genetically engineered fusion proteins and therefore avoids possible interference of tags with protein structure, interactions and function. For this CP experiment, we used highly pure mitochondria that were freshly isolated from two clones of c17orf80 knockout generated by CRISPR-Cas9 in HEK293 cells (KO), parental HEK293 cells, and parental cells treated with ddC for two or three days.

In the resulting CP dataset, 4326 proteins were identified **(Suppl. Data 3)**, of which ca. 1000 have evidence of being mitochondrial, as annotated in MitoCarta 3.0. Despite its low abundance and challenging MS detection, c17orf80 was identified (q-value: 0, score: 46.091) with 19 peptides. However, ten peptides were also detected in the KO samples. Our KO cell lines were verified to contain frameshift mutations, leading to the loss of the start codon in the third exon of c17orf80. The absence of c17orf80 protein was confirmed by Western blotting. Thus, the remaining c17orf80 peptides identified in KO samples were considered as false positives and omitted with subsequent recalculation of iBAQ values for c17orf80 based on the remaining nine identified peptides.

The signal of c17orf80 was mostly found in the high molecular mass range (>1 MDa) in two defined fractions at ∼1.3 MDa and ∼2.5-3.7 MDa. To identify potential protein interactors, we inspected the CP dataset at the position where c17orf80 was clustered **(Fig. 7A)**. Only proteins annotated in MitoCarta 3.0 were considered for this analysis. The abundance profile of c17orf80 clustered with those of a large number of proteins including RNA-binding proteins, such as mitochondrial RNA polymerases and RNA-modifying enzymes, mitoribosome-binding proteins, the large mitoribosomal subunit, as well as chaperones, proteins involved in lipid metabolism (e.g., ATAD3A), and diverse oligomeric enzymes. In addition, a portion of the replication factors TFAM and Twinkle also migrated in the same mass range, although they did not automatically cluster together with the above-mentioned proteins. This result indicates that c17orf80 exists in a high molecular mass complex with other proteins and/or nucleic acids.

**Figure 7.**
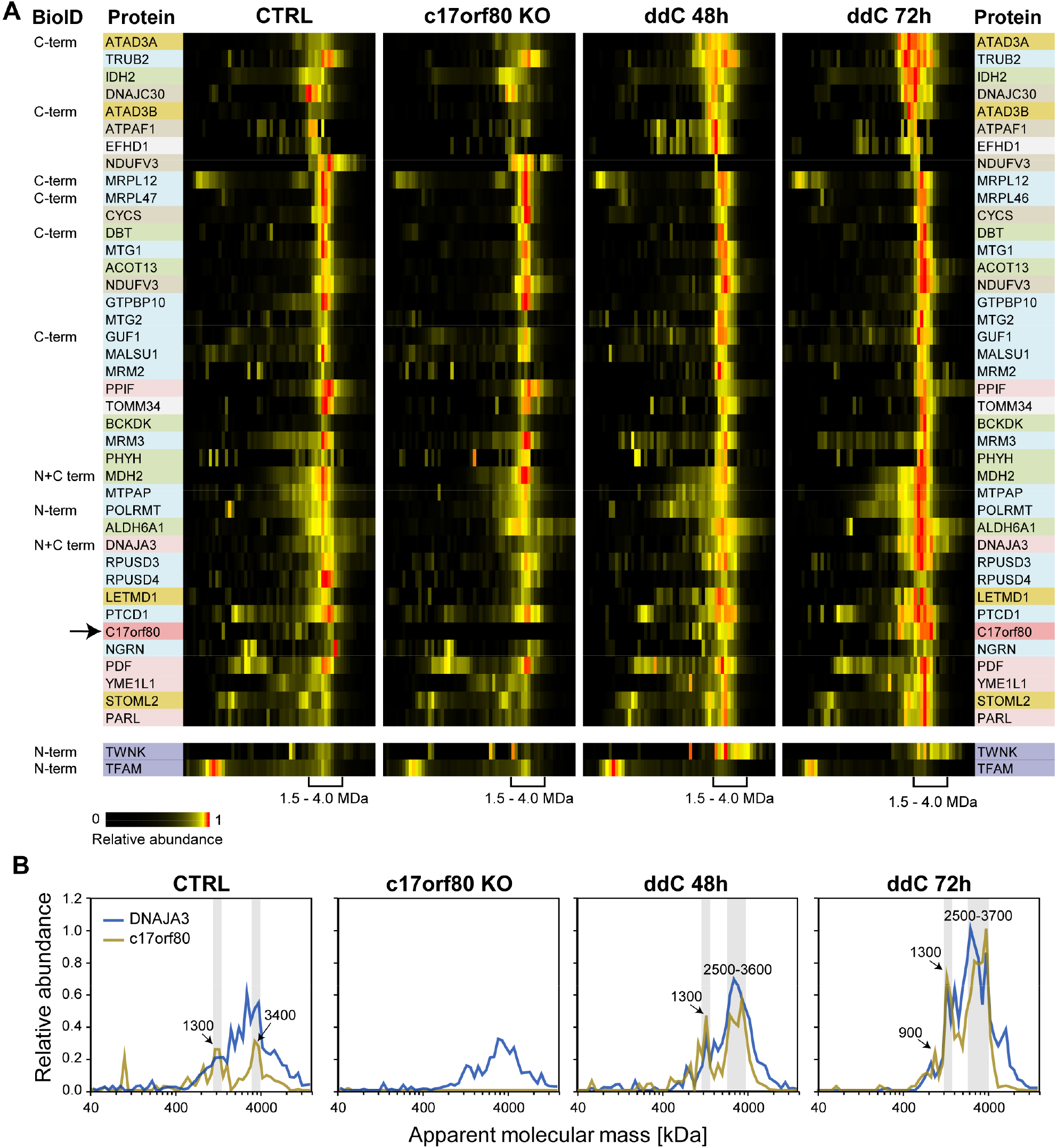
Complexome profiling of mitochondria from c17orf80 knockout and control HEK293 cells treated or not with ddC. **A**. Heatmap showing the abundance profiles of proteins co-migrating with c17orf80; low and high apparent molecular masses to the left and to the right, respectively. iBAQ values were normalized to the maximal intensity across all individual profiles (relative abundance). Corrected and averaged iBAQ values (two biological replicates of control, two KO clones, one replicate of ddC 48h and one of ddC 72h) of mitochondrial proteins were hierarchically clustered. The arrow indicates the position of c17orf80 in the cluster. Proteins are highlighted according to their main known function: RNA-binding and mitoribosome-associated proteins (blue), proteins involved in lipid metabolism (yellow), protein-modifying enzymes (pink), OXPHOS-associated proteins (grey), other oligomeric enzymes (green), mtDNA replication factors Twinkle and TFAM (purple). For visualization convenience, most of the clustered mitoribosomal proteins were excluded from the picture. Proteins enriched in C- or N-terminal c17orf80 BioID are marked. Apparent molecular masses of interest are shown as a range comprising 1.5-4.0 MDa. **B**. Abundance profiles of c17orf80 and its potential interactor DNAJA3. The y-axis represents the relative abundance. The x-axis represents the apparent molecular mass in kDa shown as log_10_-scale. The mass ranges of the major overlapping peaks are highlighted in grey.

In addition to control cells and c17orf80 KO, we analysed ddC-treated samples to determine whether the c17orf80 migration profiles change under these conditions. To evaluate the effect of ddC treatment on mitochondria, we first measured the relative mtDNA copy number and found it to decrease to 36% and 15% of the control levels after 48 h and 72 h of treatment, respectively **(Fig. S7A)**. Based on the CP data, mtDNA-encoded subunits of the OXPHOS complexes were still present at least at 40% of the control level at both time points, indicating that this residual mtDNA was sufficient to maintain gene expression active **(Fig. S7B)**. We quantified changes in protein abundance and migration using the Hausdorff effect size (*H*_*ES*_) between controls and ddC-treated samples (Hausdorff, 1927; Moeckel and Murray, 1997; Van Strien et al., 2019). In total, 421 mitochondrial proteins had different abundance profiles in ddC-treated samples based on *H*_*ES*_ > 2.0 cutoff. The most pronounced decrease was detected for the supercomplex containing OXPHOS complexes I, III and IV. Besides, we observed elevated levels of many proteins involved in mitochondrial gene expression and membrane architecture **(Suppl. Data 2)**. The abundance of c17orf80 was also higher in ddC-treated samples (*H*_*ES*_ = 1.85, 2- and 3-fold increase after 48 h and 72 h, respectively).

Among all the clustered proteins, we found that the protein DNAJA3 showed the most similar migration profile to c17orf80 **(Fig. 7B)**. Furthermore, the signal of DNAJA3 in c17orf80 KO cells was lower than in controls, whereas it was increased in ddC-treated samples like similarly to c17orf80. The abundances of DNAJA3 and c17orf80 were relatively similar in control cells, as indicated by their maximal iBAQ values of 2.6·10^6^ and 4.4·10^6^, respectively. DNAJA3 belongs to the DNAJ/Hsp40 family and acts as a co-chaperone of Hsp70 (Banerjee et al., 2022; Iosefson et al., 2012). However, the co-migration of DNAJA3 and Hsp70 was not observed in our CP dataset. Notably, we also identified DNAJA3 in our c17orf80 BioID experiment as a potential interactor. Moreover, DNAJA3 was previously found to be associated with mitochondrial nucleoids (Han et al., 2017; He et al., 2012; Lu et al., 2006; Rajala et al., 2015). Altogether, our BioID and CP results suggested an interaction between c17orf80 and DNAJA3.

### C17orf80 is not strictly required for mitochondrial gene expression in cultured cells

As we demonstrated that a substantial portion of c17orf80 resides in mitochondrial nucleoids, we aimed to investigate possible effects of its loss on mtDNA maintenance and gene expression. To this end, we knocked out c17orf80 in HEK293 cells using CRISPR-Cas9 system. We generated two single-cell clones of c17orf80 knockout (KO-1 and KO-2) and compared them with the parental cells **(Fig. 8A)**.

**Figure 8.**
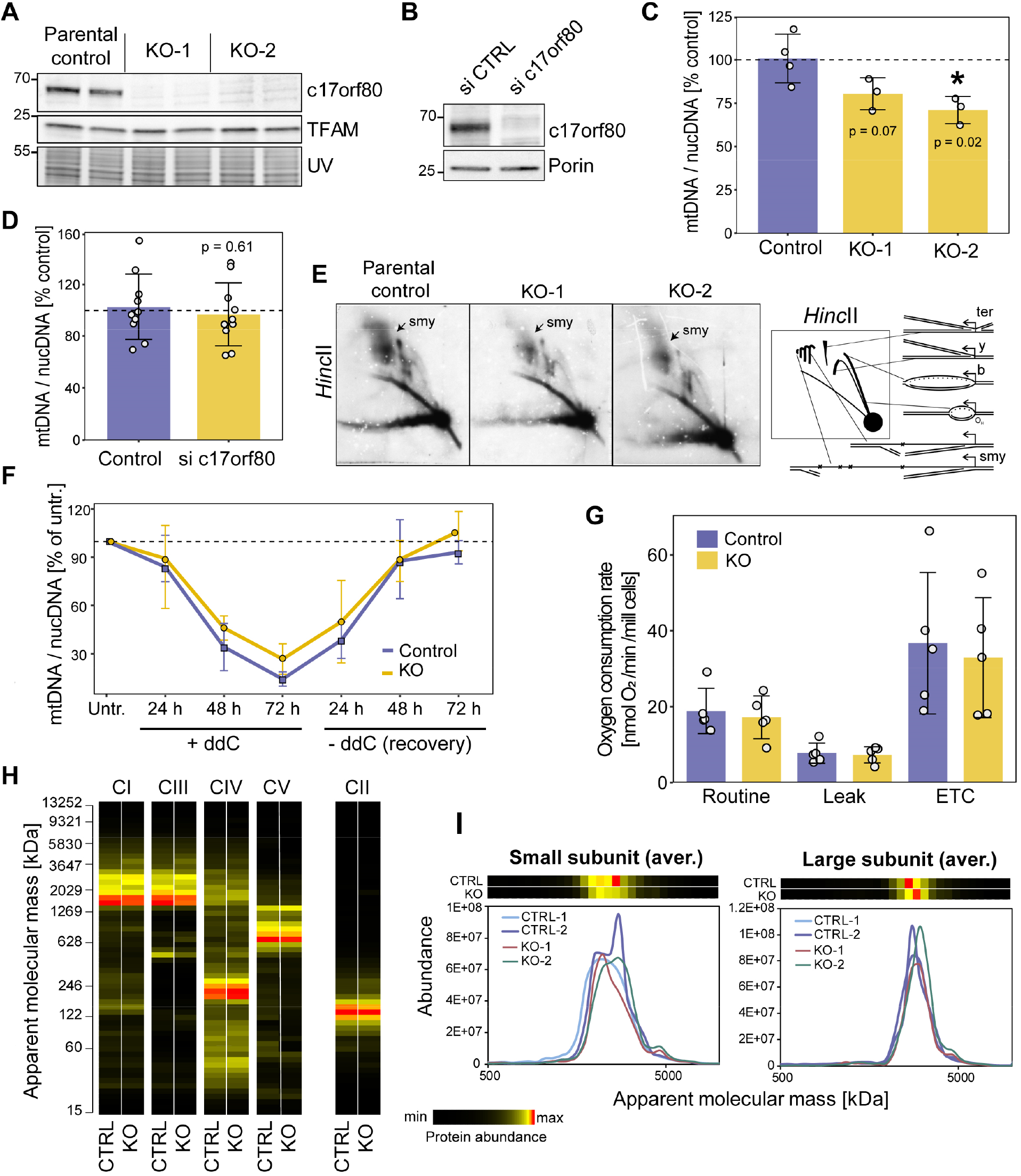
Lack of c17orf80 does not have a major effect on mtDNA copy number, replication and mitochondrial gene expression. **A**. Levels of c17orf80 and TFAM in parental HEK293 cells and c17orf80 KO clones. Protein staining detected under UV light was used as a loading control. **B**. Levels of c17orf80 in control cells and siRNA-mediated knockdown three days after transfection. Porin was used as a loading control. **C-D**. mtDNA content was not significantly affected by depletion of c17orf80. qPCR quantification of relative mtDNA copy number in KO (C) and knockdown (D). Data are mean ± SD from three or four independent measurements for KO and ten independent knockdowns; individual values are overlaid; unpaired Student’s t-test. **E**. mtDNA replication intermediates in KO and control cells analysed by 2D-AGE of mtDNA digested with *Hinc*II. On the left, a schematic representation of the replication intermediates is shown: Ter -termination intermediates; y – y-arc; b – bubble arc; smy – slow moving y-arcs. **F**. KO and control cells behave similarly during ddC treatment-recovery time course. Cells were treated with 100 μM ddC for 24-72 hours and allowed to recover in the ddC-free medium for 24-72 h. Relative amount of mtDNA was measured at every time point as a ratio of CytB to APP genes using qPCR. For each experiment, the value measured for cells cultured without the addition of ddC was set to 100%. Data are means ± SD from three independent experiments. **G**. Mitochondrial OXPHOS function was not affected in c17orf80 KO. OCR values from routine, leak and ETC were corrected to CS activity (U/g). Data are mean values ± SD from five independent experiments with overlaid individual values; unpaired Student’s t-test. **H-I**. Complexome profiling of KO and control cells. The heatmaps and plots represent averaged intensities of all detected individual subunits for each OXPHOS complex (I) and mitoribosome subunits (J) from two KO clones and two biological replicates of control. Complexes I, III, IV and V are labelled as CI, CIII, CIV and CV, respectively. Complex II (CII) is shown as loading control as it does not contain mtDNA-encoded subunits.

First, we measured the relative mtDNA copy number in c17orf80 KO cells using real-time PCR (quantitative PCR, qPCR). We found that both clones had a moderate reduction in mtDNA content (a 20-30% decrease compared to parental control), which however was statistically significant only for one of the clones **(Fig. 8C)**. Since metabolic compensation might occur in knockout cells masking an mtDNA phenotype, we also checked whether the mtDNA content was affected by a short-term depletion of c17orf80. For this, we transfected U2OS cells with small interfering RNAs (siRNAs) targeting c17ofr80 or with a negative siRNA control **(Fig. 8B)** and measured the relative mtDNA copy number in ten independent biological replicates of knockdown and control. We found no change in mtDNA content after three days of c17orf80 depletion **(Fig. 8D)**, suggesting that c17orf80 is not directly involved in mtDNA copy number regulation.

Next, we investigated whether the lack of c17orf80 caused any mtDNA maintenance perturbations. We examined mtDNA replication intermediates in c17orf80 KO cells and parental control cells using 2D DNA agarose electrophoresis **(Fig. 8E)**. We observed a slight decrease of the slow-moving replication forks (y-arcs), containing intermediates of the RITOLS mtDNA replication (Pohjoismaki et al., 2010; Yasukawa et al., 2006), in both c17orf80 KO clones. There was no mtDNA replication stalling in response to c17orf80 absence. We also measured levels of 7S DNA using Southern blotting and found no difference between c17orf80 KO and parental control cells **(Fig. S8A)**. In addition, we analysed the topology of mtDNA using Southern blotting. We observed that KO cells exhibited a minor but significant increase in oligomeric and open circle forms of mtDNA accompanied by a slight decrease in supercoiled species and replicating mtDNA molecules **(Fig. S8B)**. Overall, the changes in mtDNA replication mode and topology were minor and require further validation.

Furthermore, we assessed the morphology of mitochondria and nucleoids using immunofluorescence microscopy and found no visible changes upon c17orf80 silencing in U2OS cells **(Fig. S9)**.

Since we previously observed an accumulation of c17orf80 in nucleoids upon ddC treatment, we decided to test the effect of ddC exposure on mtDNA depletion and the efficiency of mtDNA recovery after its removal in KO and control cells. We measured the relative mtDNA copy number every 24 h throughout the three days of ddC treatment and the three days after its removal. We did not observe any substantial changes in the rates of mtDNA depletion and recovery between control and c17orf80 KO **(Fig. 8F)**.

Due to its association with mtDNA and mitochondrial gene expression factors, we tested whether the lack of c17orf80 affects the function of the OXPHOS system. To this end, we analysed the mitochondrial respiration in c17orf80 KO cells. We measured the oxygen consumption rates (OCRs) in freshly harvested intact cells using an Oroboros™ Oxygraph-2k. We found no difference in the OCRs of the routine, leak, and maximal respiration states between c17orf80 KO and control cells **(Fig. 8G)**.

To determine whether c17orf80 KO cells carry any defects in mitochondrial protein complexes architecture, we used our CP data of c17orf80 KO for detailed examination of OXPHOS, mitoribosome and other proteins. As expected, KO cells showed no sign of OXPHOS changes as the subunits of all four complexes containing mtDNA-encoded subunits, i.e. complexes I, III, IV and F_1_F_O_-ATP synthase (CV), and their respective supercomplexes (Signes and Fernandez-Vizarra, 2018) were properly assembled in amounts comparable to control **(Fig. 8H)**. No changes were observed for the mitoribosome **(Fig. 8I)**. This suggests that c17orf80 does not relate to biogenesis of OXPHOS complexes and its absence does not dramatically affect the expression level of mitochondrial transcripts. In addition, we quantified changes in the protein abundance profiles between control and KO using the Hausdorff effect size for all 4326 identified proteins. We found no systematic changes between the complexomes of c17orf80 KO and control cells **(Suppl. Data 3)**.

Taken together, these data indicated that c17orf80 is not strictly required for mtDNA replication and gene expression in human cells grown under normal culturing conditions.

## Discussion

Thousands of human proteins remain poorly characterized despite the reported involvement in crucial cellular processes and diseases (Haynes et al., 2018; Kustatscher et al., 2022; Rensvold et al., 2022; Sinha et al., 2018; Stoeger et al., 2018). Proper execution of the mitochondrial gene expression system is essential for a healthy energy metabolism. In this study, we investigated an uncharacterized protein c17orf80, which was previously captured by mitochondria-targeted proximity mass spectrometry. Using immunofluorescence microscopy and interaction proteomics, we demonstrated that c17orf80 associates with the mitochondrial nucleoid, a nucleoprotein complex composed of mtDNA and proteins involved in its maintenance, replication and transcription. In addition, we showed that although c17orf80 interacts with the nucleoids, its loss did not lead to a mitochondrial dysfunction in human cells cultured under standard growth conditions.

Here, we provide evidence that c17orf80 is a novel nucleoid resident, as evaluated by both immunofluorescence microscopy and BioID analysis. Notably, c17orf80 was detected only in recent proximity-labelling proteomic experiments (Antonicka et al., 2020; Hangas et al., 2022; Hensen et al., 2019; Jiang et al., 2019), but was not identified in early IP-based studies or biochemical nucleoid preparations (Bogenhagen et al., 2008; He et al., 2012; Rajala et al., 2015). This probably reflects the fact that biotin-based proximity labelling techniques allow better detection of labile protein interactions that potentially do not withstand standard affinity-purification procedures. Our IF experiments showed that c17orf80 forms a punctate pattern colocalizing with mtDNA, which is a characteristic feature of core nucleoid proteins such as TFAM, Twinkle, and mtSSB (Spelbrink, 2010). Furthermore, c17orf80 retained in nucleoids under replication stress caused by ddC and EtBr, suggesting it has a high affinity to one of the nucleoid components, either mtDNA or a protein. These findings were supported by the BioID proximity-biotinylation experiment that showed that c17orf80 is located in close spatial proximity to mitochondrial replication and gene expression factors.

The secondary structure of c17orf80 mostly lacks defined functional motifs, hampering any hypothesis about its molecular function. There is, however, a possible CCHC-type ZnF located on its N-terminus suggesting nucleic-acid binding abilities. C17orf80 may acquire more structural features upon binding to another protein or nucleic acids, as it is often true for intrinsically disordered proteins (van der Lee et al., 2014; Wright and Dyson, 2009). In various organisms, some proteins with such highly disordered sequences are involved in DNA damage protection (Hashimoto and Kunieda, 2017; Smith and Graether, 2022) and repair (Balasubramanian et al., 2022; López et al., 2020) via direct interaction with DNA.

We observed c17orf80 to pellet with the membrane fraction after sodium carbonate extraction, which is consistent with the structural prediction of two TM helices at its C-terminus. Since a portion of c17orf80 was also present in the supernatant, resembling peripheral membrane proteins (Kim et al., 2015), we propose that it peripherally associates with the IMM with the majority of the polypeptide being in the mitochondrial matrix. Based on the results of our antibody-accessibility assay and BioID, we excluded a stable association with the OMM. Additionally, we detected homology of the C-terminus of c17orf80 to the subunit f of the F_1_F_O_-ATP synthase (CV). However, we found no experimental support for an interaction of this protein with CV.

Interestingly, although the core mtDNA-binding components of nucleoids are soluble replication and transcription factors, nucleoids have been proposed to interact with the inner mitochondrial membrane (Hillar et al., 1979), possibly via its peripheral components (Bogenhagen et al., 2008; Gerhold et al., 2015; He et al., 2012; Kasashima et al., 2008). In addition, a recent super-resolution microscopy study showed that most of mtDNA colocalizes with a membrane-scaffolding protein ATAD3A (Arguello et al., 2021), supporting this model. In yeast, tethering nucleoids to the inner membrane is essential for mtDNA integrity (Dimmer et al., 2005). Considering that we did not observe enrichment of ATAD3A in the c17orf80 interactome or any changes in nucleoid morphology or mtDNA depletion upon loss of c17orf80, it is unlikely that this protein is a major player in nucleoid membrane attachment.

Previously, the interactome of c17orf80 was investigated by Antonicka et al., using c17orf80 C-terminally tagged with the biotin ligase (Antonicka et al., 2020). Following the analysis of 100 BioID baits, the authors assigned c17orf80 to a cluster of RNA-processing factors located in mitochondrial RNA granules. It is common to use C-terminal tags for mitochondrial proteins to prevent disruption of the mitochondria-targeting sequence, which is usually located at the N-terminus. However, in the case of c17orf80, we showed that the N-terminal fusion with BirA* was successfully imported into mitochondria and labelled a distinct set of proteins compared to the C-terminal version. Using this approach, we identified TFAM, mtSSB and Twinkle as the c17orf80 interactors, confirming the nucleoid association of c17orf80. We did not identify many RNA granule residents in this experiment and did not detect colocalization of c17orf80 with BrU-labelled RNA-granules by IF. Thus, possible involvement in RNA processing was not investigated here; however, it cannot be excluded with our current evidence.

Based on our BioID and complexome profiling results, we suggested an interaction between c17orf80 and DNAJA3 (a.k.a. Tid1), a member of the DNAJ/Hsp40 protein family and co-chaperone of Hsp70 (Banerjee et al., 2022; Syken et al., 1999). DNAJA3 has been shown to be associated with nucleoids in several studies (Han et al., 2017; He et al., 2012; Lu et al., 2006; Rajala et al., 2015), and its depletion was linked to reduced mtDNA copy number and OXPHOS deficiency (Hayashi et al., 2006; Ng et al., 2014; Wang et al., 2020). It was suggested that these effects may be linked to DNAJA3 co-chaperone activity and accumulation of aggregated complex I upon its depletion (Ng et al., 2014). The direct involvement of DNAJA3 in mtDNA-related processes is still to be investigated. It is necessary to note that the quantification of both c17orf80 and DNAJA3 by CP was challenging, with relatively low intensity, some misidentified and low-scoring peptides, and dispersed profiles at the high molecular mass range. However, since our CP data are based on four independent profiles, where the levels of the proteins correlate with each other and are supported by the identification of DNAJA3 in c17orf80 BioID, the interaction of these two proteins is plausible and deserves to be investigated in more detail.

We observed a prominent accumulation of c17orf80 in nucleoids upon treatment with ddC, a modified cytosine analogue that causes stalling of mtDNA replication via inhibition of Polγ processivity and/or nascent strand termination (Brown and Clayton, 2002; Nelson et al., 1997). It is known that the rapid mtDNA depletion induced by ddC is accompanied by complete inhibition of mtDNA replication (Rajala et al., 2014) and TFAM degradation (Li et al., 2021). However, the molecular mechanisms of subsequent mtDNA decay and its repair have not been extensively investigated. Several proteins have been proposed to participate in mtDNA repair or restart of mtDNA replication under or after ddC exposure, such as PrimPol (Carvalho et al., 2021), MGME1 (Kornblum et al., 2013), RAD51C/XRCC3 complex (Mishra et al., 2018), and TDP1 (Huang et al., 2013). Generally, in the absence of these proteins, cells were unable to recover mtDNA levels after its depletion. In our c17orf80-lacking cells, mtDNA depletion and recovery rates were similar to those in the control, suggesting that the accumulation of c17orf80 in ddC-treated nucleoids is not a sign of its direct involvement in mtDNA repair or replication, at least not in HEK293 cells.

In our CP experiment performed with ddC-treated cells, a three days exposure to ddC did not lead to complete depletion of OXPHOS complexes and their mtDNA-encoded subunits, even when the mtDNA level was diminished to 15% of the control after three days. Instead, we observed upregulation of transcription and translation factors, mitoribosomes, and complexes involved in cristae formation (e.g. ATAD3A, MICOS complex) and protein quality control (e.g. ClpX), likely reflecting the general compensatory response to mtDNA depletion. C17orf80 was also elevated in ddC-treated samples, which correlates with our IF experiments demonstrating its enrichment in nucleoids after ddC exposure.

Despite its nucleoid localization, c17orf80-deficient cells showed no major signs of mtDNA replication impairment or changes in nucleoid morphology. We also did not detect gene expression defects, as OXPHOS complexes I, III, IV and V were fully assembled and functioning in KO cells.

In summary, we identified c17orf80 as a novel membrane-associated resident of mitochondrial nucleoids. It is important to note that it is possible that the molecular function of c17orf80 is not relevant for the type of cells and growth conditions used in this study. In such manner, c17orf80 may be more active only, for instance, under specific stress conditions, such as rapid mtDNA depletion, and may be involved in a molecular process related to mtDNA protection or compensatory gene expression. Further studies are certainly required to define the specific molecular role of c17orf80 in nucleoids from other cell types and tissues, especially under stress caused by mtDNA replication inhibitors.

## Methods

### Sequence analysis

The homology of the C-terminus with the F_1_F_O_-ATP synthase subunit f and the low level of overall sequence conservation of c17orf80 hampers large scale automatic orthology detection. We therefore combined the relatively strictly defined orthologs of Ensembl from c17orf80 in human and those of the *Danio rerio*’s c17orf80 (ENSDARP00000110900), and an alignment was created with clustalx (Sievers and Higgins, 2018). This alignment was used to create the sequence conservation using ConSurf (Ashkenazy et al., 2016) and sequence logo with WebLogo (Crooks et al., 2004) **(Fig. 1C and 1D)**, and to run an HHpred search (Zimmermann et al., 2018) (0 iterations) against PDB and PFAM databases **(Fig. S1A)**, while the alignment of the sequence logos of the ATP synthase subunit f and the c17orf80 C-terminal domain **(Fig. S1B)** was created with HHpred, using alignments of the vertebrate ATP synthase subunits f and the c17orf80 C-terminal domain.

### Cell culture and treatments

HEK293 (ATCC; CRL-1573), Flp-In T-REx 293 (Invitrogen), and U2OS cells (University of Helsinki, Finland) were grown in high-glucose DMEM (Lonza; BE12–604F) supplemented with 10% fetal bovine serum (FBS) (Gibco; 16000044) in a humidified 37 °C incubator at 5% CO2. All cell lines were routinely tested for mycoplasma contamination and found to be negative. The Flp-In T-Rex cells were grown with the addition of blasticidin or zeocin, and hygromycin according to the manufacturer’s protocol. To inhibit mtDNA replication, growth medium was supplemented with 100 μM of 2′,3′-dideoxycytidine (Sigma; D5782) or 50 ng/ml of ethidium bromide. For the detection of RNA granules, cells were incubated with 2.5 mM of BrU (Sigma; 850187) for 20 or 60 min before immunofluorescence detection.

### Immunofluorescence

For immunofluorescence detection, U2OS cells were grown on coverslips in six-well plates. The cells were fixed using 3.3% paraformaldehyde (PFA) in DMEM/10% FBS for 20 min, washed in phosphate-buffered saline (PBS), and permeabilized for 15 min with 0.5% Triton X-100 (TX-100) in PBS/10% FBS. The cells were then incubated for 1 h with primary antibodies diluted in PBS/10% FBS (Table 1). After washing, the cells were incubated for 45 min with the corresponding secondary antibodies labelled with Alexa Fluor diluted in PBS/10% FBS (1:1000; Invitrogen): goat-anti-rabbit IgG 488 nm, goat-anti-mouse IgG 568 nm, and goat-anti-mouse IgM 647 nm. After the final incubation and washing, coverslips were mounted using ProLong™ Gold Antifade reagent with DAPI (Invitrogen; P36935). Images were acquired using a Zeiss apotome system in apotome mode on an Axio Observer Z.1 with Colibri-led illumination and appropriate emission filters using the x40 or x63 oil immersion objectives. Images were processed using the ImageJ software (Schneider et al., 2012). To compare control cells with cells treated with chemicals or siRNA, images for each channel were acquired with identical illumination and exposure settings and processed identically.

**Table 1.** Antibodies used for IF detection

|  |  |  |  |  |
| --- | --- | --- | --- | --- |
| Anti-c17orf80 | Rabbit IgG | Proteintech | 27762-1-AP | 1:200 |
| Anti-TFAM | Mouse IgG | Abcam | ab89818 | 1:200 |
| Anti-Cyclophilin D | Mouse IgG | Abcam | ab110324 | 1:400 |
| Anti-BrU | Mouse IgG | Roche | 11170376001 | 1:50 |
| Anti-DNA | Mouse IgM | Progen | 61014 | 1:100 |
| Anti-Tom20 | Mouse IgG | Santa Cruz | sc17764 | 1:100 |
| Anti-MRPL12 | Mouse IgG | Abcam | ab58334 | 1:400 |
| Anti-mtSSB | Rabbit IgG | Sigma | SAB4502859 | 1:200 |
| Anti-POLRMT | Rabbit IgG | Abcam | ab32988 | 1:100 |
| Anti-Twinkle | Mouse IgG | gift from Anu Suomalainen Wartiovaara |  | 1:50 |

### Antibody accessibility assay

For the immunofluorescent antibody accessibility assay, PFA-fixed cells were treated with 100 μM digitonin (Millipore; 300410) diluted in PBS for 7 min at room temperature, washed with PBS and either or not further permeabilized with 0.5% TX-100 for 15 min, followed by antibody incubation as described above.

### Image quantification

For colocalization analysis, Manders colocalization coefficients M1 and M2 were calculated using the JaCop plug-in (Bolte and Cordelieres, 2006) in ImageJ with automatically selected intensity thresholds. For intensity and particle size quantification, the processed TIF files for each channel were subjected to an intensity threshold and watershed separation. The obtained particles were quantified using basic ImageJ quantification functions. Summary data of the mean grey values and average particle sizes from each image were used for statistical analysis. To analyse only mitochondria-derived DNA signal, the area of the cell nuclei was manually cut out before colocalization and particle analysis for all three channels.

### Isolation of mitochondria

Mitochondria from HEK293 cells were isolated as follows: cells were harvested by resuspension in growth medium, washed with cold PBS, pelleted, and resuspended in isotonic isolation buffer (250 mM sucrose, 1 mM EDTA, 20 mM Tris-HCl, pH 7.4, protease inhibitor cocktail SigmaFAST). Cells were disrupted by 10-15 strokes in a Potter-Elvehjem tissue grinder. The homogenate was diluted with three volumes of isolation buffer and centrifuged (1000 g; 10 min; 4 °C) to remove the debris and cell nuclei. The crude mitochondrial fraction was pelleted from the cleared supernatant (11 000 g; 10 min; 4 °C). For complexome profiling, the mitochondrial pellet was re-suspended in 0.5 ml of isolation buffer, loaded on a two-step sucrose gradient (1.5 M/1 M sucrose in 20 mM Tris-HCl pH 7.4, 1 mM EDTA), and centrifuged at 60 000 g for 20 min at 4 °C in a swing-out rotor. The highly pure mitochondrial fraction was recovered from the interphase between sucrose layers and washed with three volumes of isolation buffer (11 000 g; 5 min; 4 °C). All the above steps were performed on ice using ice-cold buffers. Protein concentration was determined using the Bradford colorimetric assay (Quick Start™ Bradford Protein Assay Kit; Bio-Rad; 5000201).

### Sodium carbonate extraction

Mitochondrial pellets containing 200 μg of proteins were re-suspended in either 400 μl of the isolation buffer or 400 μl of freshly prepared 100 mM sodium carbonate (pH 11.0) and incubated on ice for 30 min. The mitochondrial membranes were pelleted at 60 000 g for 40 min at 4 °C. The supernatants were transferred to new tubes, and the pellets were carefully washed with the isolation buffer or sodium carbonate. Proteins from supernatants were precipitated with 20% TCA for 15 min on ice, pelleted (18 000 g; 10 min; 4 °C) and washed twice with ice-cold acetone (18 000 g; 3 min; 4 °C). The protein pellets were resuspended in Laemmli sample buffer and subjected to SDS-PAGE and Western blotting. For quantification, western blot bands were quantified in Image Lab 4.1 Software (Bio-Rad) using volume tools.

### Generation of c17orf80-BirA* stable cell lines

Plasmids encoding for c17orf80-BirA* N- or C-terminal fusions were generated using Gateway cloning technology (Invitrogen) using the c17orf80 sequence (NM_001100621) amplified from a commercially available tagged ORF clone (OriGene; RC221916L3) and pDEST-pcDNA5-BirA-FLAG vectors (Couzens et al., 2013).

Flp-In T-REx 293 cells were grown in antibiotic-free media on 10 cm dishes to 80% confluency. Cells were co-transfected with 3.6 μg of pOG44 and 400 ng of either N- or C-terminal pDEST-pcDNA5-c17orf80-BirA*-FLAG using Lipofectamine 2000 (Invitrogen; 11668019) according to the manufacturer’s protocol. The next day, antibiotics were added to the transfected cells to allow for clonal selection (hygromycin B 200 μg/ml; blasticidin 7 μg/ml). After two days, detached cells were removed and medium with lower antibiotics concentrations was added (hygromycin B 100 μg/ml, blasticidin 7 μg/ml). This medium was refreshed every 3-4 days until hygromycin-resistant colonies formed. After formation, the colonies were transferred to new dishes, expanded and frozen for storage. Cell lines expressing the fusion constructs were tested for optimal concentration of the inducing agent (anhydrotetracycline, AnTET).

### BioID protein interactome mapping

The BioID pulldowns were performed generally as described previously (Antonicka et al., 2020). Flp-In T-REx cell lines expressing c17orf80-BirA* fusions were grown on 15 cm dishes without antibiotics for two days they until reaching 70% confluency. To induce fusion protein expression, AnTET was added to a final concentration of 1 ng/ml (N-term) or 3 ng/ml (C-term). Simultaneously, biotin was added at a final concentration of 50 mM. For control pulldowns, the same cell lines were supplemented with biotin alone and without AnTET. Cells were harvested 22-24 hours after biotin supplementation. Equal amounts of cells (3·10^7^) were snap-frozen and stored at -80 °C until affinity purification.

For biotin-streptavidin pulldowns, cell pellets from four biological replicates were thawed on ice and lysed with 2 ml of lysis buffer (50 mM Tris-HCl pH 7.5, 150 mM NaCl, 1% NP-40, 1 mM EDTA, 0.1% SDS, protease inhibitors, 0.5% sodium deoxycholate) for 1 h on a roller shaker at 4 °C. The lysates were sonicated on ice with a microtip sonicator (120 W, 8 s on, 2 s off, 3 pulses) and supplemented with 200 U of benzonase (Sigma; E8263). The lysates were cleared by centrifugation (18000 g; 30 min; 4 °C). The recovered supernatants were incubated with 50 μl of Pierce™ streptavidin agarose beads (Thermo Fisher Scientific; 20353), pre-equilibrated in washing buffer, overnight on a roller shaker at 4 °C. The next day, beads were collected (400 g; 5 min; 4 °C) and washed as follows: twice with 2 ml of washing buffer (50 mM Tris-HCl pH 7.5, 150 mM NaCl, 1% NP-40, 1 mM EDTA, 0.1% SDS); twice with 2 ml of 8 M urea freshly prepared in 10 mM Tris-HCl, pH 7.5; thrice with 1.5 ml of 50 μM ammonium bicarbonate (ABC) prepared with HPLC-grade water. All steps were performed on ice using ice-cold buffers.

After the final wash, the beads were transferred to new 1.5 ml Eppendorf Protein LoBind Tubes and subjected to reduction with 50 μl 2 mM dithiothreitol (30 min; 25 °C; shaking), alkylation with 50 μl of 10 mM chloroacetamide (30 min; 25 °C; shaking), and digestion with 0.5 μg trypsin (Promega; V5111) prepared in 200 μl 50 mM ammonium bicarbonate (ABC) (overnight; 37 °C; horizontal shaking). After digestion, the beads were collected (800 g; 2 min) and washed twice with 50 mM ABC to gather the remaining peptides (150 μl; 800 g; 2 min). The supernatants after digestion and washes were combined and dried in a vacuum concentrator. The dried peptides were resuspended in 20 μl of 5% acetonitrile (ACN) in 0.5% formic acid (FA) and analysed by LC-MS/MS as described previously (Evers et al., 2021) with minor modifications. Here, peptides were separated on a 60 min linear gradient of 5-35% ACN/0.1% FA, followed by 35-80% gradient of ACN/0.1% FA (5 min) at a flow rate of 300 nl/min, and a final column wash with 90% ACN (5 min) at 600 nl/min.

MaxQuant v.1.6.10.43 was used to match the identified tryptic peptides to the human UniProt database (ID UP000005640, release date 28/04/2021). Label-free quantification (LFQ) was used to quantify the identified proteins. Data analysis was performed using Perseus v.1.6.15.0 (Tyanova et al., 2016). Only proteins identified in at least three biological replicates of c17orf80-BirA*(N) or -BirA*(C) pulldowns were included in the analysis. LFQ values were log_2_-transformed and missing values were replaced by random values drawn from a normal distribution (width = 0.3, downshift = 1.8). The log_2_ fold change values (log_2_FC) were calculated as the difference between log_2_-transformed averaged quadruplicates. An unequal variance unpaired t-test (Welch’s) was used to test for the null hypothesis of no difference between protein abundances. Statistical significance was determined using a permutation-based FDR < 5% threshold with artificial within-group variance S0 = 0.1 (Tusher et al., 2001). For data visualization, negative log_10_-transformed *p* values were plotted against log_2_FC in R-studio.

### Transient protein depletion by siRNA

For c17orf80 knockdown, U2OS cells were transfected with a mixture of three Stealth™ siRNA duplex oligonucleotides (Invitrogen; HSS123763, HSS123765, HSS182809) at a concentration of 10 nM each, using Lipofectamine 2000 (Invitrogen; 11668019), according to the manufacturer’s protocol. The cell culture medium was changed 4-6 hours after transfection. Cells were analysed 68-72 hours after transfection. Stealth RNAi™ siRNA Negative Control mix (Invitrogen; 12935300) was used for control transfections.

### CRISPR-Cas9 knockout

The guide RNA directed against the third exon of c17orf80 was designed using the available online tools with the following sequence: GCTGGAGCGTCTTTACTGGT**TGG**. The gRNA was inserted into the pSpCas9(BB)-2A-GFP vector (PX458; Addgene #48138) following a previously published protocol (Ran et al., 2013). For nucleofection, 1·10^6^ of HEK293 cells were resuspended in 100 μL of pre-warmed Nucleofector solution V (Lonza). The plasmid containing c17orf80 gRNA and Cas9 (4 μg) was added to the cell suspension. The suspension was transferred to an electroporation cuvette and electroporated using the Q-001 program on an Amaxa Nucleofector II (Lonza). The cells were recovered from the cuvette, seeded in a 6-well plate, and allowed to recover for one day. Next, the cells were harvested, counted, and seeded as single cells in 96-wells using a serial dilution method. In approximately a week, the wells containing single-cell clones were identified and marked. After single-cell clones formed colonies, they were transferred to 24-wells plates, expanded and screened for the absence of c17orf80 protein expression by SDS-PAGE and Western blotting with antibodies against c17orf80. For clones in which no c17orf80 protein was detected, the third exon was analysed using Sanger sequencing to confirm gene editing.

### mtDNA copy number measurement

Total DNA from cultured cells was isolated using a NucleoSpin Tissue Purification Kit (Macherey-Nagel; 740952) according to the manufacturer’s protocol. The mtDNA content was quantified and normalized to nuclear DNA (nucDNA). Measurements were performed using quantitative real-time PCR (qPCR) with primers for cytochrome *b* (CytB, mtDNA) and amyloid precursor protein (APP, nucDNA): CytB-Fw GCCTGCCTGATCCTCCAAAT, CytB-Rv AAGGTAGCGGATGATTCAGCC; APP-Fw TTTTTGTGTGCTCTCCCAGGTCT, APP-Rv TGGTCACTGGTTGGTTGGC. Each qPCR reaction contained 25 ng of purified total DNA, 2.5 mM of forward and reverse primers, 10 μl of 2× SYBR Green Master Mix (Bio-Rad; #4309155) and was measured in triplicate in Hard-Shell 96-Well PCR Plates (Bio-Rad; #HSP9635). The amplification program was as follows: 95 °C for 10 min followed by 40 cycles of 95 °C for 15 s and 60 °C for 60 s. The fluorescent signal was acquired by CFX96 Real-Time System (Bio-Rad). The absence of non-specific amplicons was confirmed using melting curve analysis. Fold changes in the relative mtDNA copy number after c17orf80 depletion or ddC treatment were calculated using the 2^−ΔΔCT^ method.

### 2D agarose gel electrophoresis

Two-dimensional agarose gel electrophoresis (2D-AGE) was performed as described previously (Hangas et al., 2022). Briefly, 5 μg of mtDNA was digested with *Hinc*II, separated by 2D-AGE and blotted. The membranes were probed with a ^32^P-labelled probe (nucleotides 37–611 of human mtDNA) and exposed on Kodak Biomax MS film using an intensifying screen.

### mtDNA topology analysis

Analysis of topological forms of mtDNA was performed as described previously (Hangas et al., 2022). Briefly, 1.5 μg of total DNA digested with *Bgl*II was separated over a 0.4% agarose gel in TBE, blotted and probed with a ^32^P-labelled probe against nucleotides 37-611 of human mtDNA. The blot was exposed and the various bands quantified by phosphor imaging.

### 7S DNA quantification

Quantification of mitochondrial 7S DNA levels per mtDNA was performed as described previously (Hangas et al., 2022). Briefly, 1.5 μg of total DNA was digested with *BamHI*, heated for 10 min at 65°C, separated over a 0,8% agarose gel in TAE, blotted and probed with a ^32^P-labelled probe against nucleotides 16,177-40 of human mtDNA (7S). 7S DNA and full-length mtDNA were quantified using a phosphor imager and the ratio of 7S per mtDNA band intensities was calculated.

### Complexome profiling

Freshly obtained mitochondrial pellets (200 μg proteins) were solubilized with digitonin (8 g/g protein; SERVA) in hypotonic buffer (50 mM NaCl, 2 mM aminohexanoic acid, 50 mM imidazole, 2.5 mM MgCl_2_, 2 mM CaCl_2_, pH 7.0) supplemented with 150 U of DNase I (Invitrogen; 18047019) and kept at +10 °C for one hour. Lysates were cleared by centrifugation (22000 g; 20 min; 4 °C). The supernatants were recovered, supplemented with 2.5 mM EDTA and Ponceau loading buffer, and separated on a 3-16% high-resolution clear native polyacrylamide gel (hrCN-PAGE) for 30 min at 100 V, followed by 4 h at 200 V and 20 min at 400 V. It should be noted that DNase I was added during the solubilization to let DNA-binding proteins enter the native gel. Since all samples were supplemented with MgCl_2_ for effective DNA digestion, hrCN-PAGE was used here to avoid precipitation of Coomassie blue dye and proteins by divalent cations as it may happen when using widespread blue native (BN)-PAGE (Wittig et al., 2006). After the run, the gel was processed as described previously (Evers et al., 2021) and each line was cut into 56 slices. In-gel digestion, mass spectrometry analysis and complexome profiling were performed as described previously (Evers et al., 2021) with some modifications. Here, MaxQuant v.2.0.3 was used to match identified tryptic peptides to the human UniProt database (ID UP000005640, release date: 28/04/2021). Data were normalized for the sum of iBAQ values of mitochondrial proteins annotated in MitoCarta 3.0 from each sample. The mass calibration was performed using human membrane and soluble protein complexes of known masses (e.g., OXPHOS, VDACs, HSP60, OGDH; see **Suppl. Data 3**). To correct for differences between multiple gel runs, the profiles were aligned using the COPAL tool (Van Strien et al., 2019). In addition, Hausdorff effect sizes (*H*_*ES*_) were calculated in COPAL between control (n = 2) and c17orf80 KO (n = 2) and between control (n = 2) and ddC (n = 2) samples to evaluate the difference between protein migration profiles. Proteins with *H*_*ES*_ > 2.0 were generally considered to be affected. Prior to averaging the abundance profiles from the proteins of interest, intensity signals that were not identified in both replicates were manually excluded.

### High-resolution respirometry

The oxygen consumption rates (OCRs) in intact cells in suspension were measured using a high-resolution respirometry system (Oroboros™ Oxygraph-2k). Control and c17orf80 KO cells were harvested, counted and placed in 2 ml chambers filled with DMEM/10% FBS at a concentration of 1·10^6^ cells/mL at 37 °C with stirring at 750 rpm. First, the basal respiration representing the endogenous coupled respiration state was recorded (Routine). Oligomycin (1 μg/ml) was added to the chambers to inhibit mitochondrial ATP synthesis and the non-phosphorylating respiration was measured (Leak). CCCP was then added stepwise, first adding 2.5 μM shots, followed by several 1 μM shots until the maximal uncoupled respiration (ETC) was reached. Finally, mitochondrial respiration was inhibited by adding 30 μM rotenone and 2.5 μM antimycin A. Residual non-mitochondrial respiration was recorded for 10 min. To normalize the OCRs, the activity of citrate synthase (CS) and protein concentration were measured in each sample using a KoneLab 20XT platform, as described previously (Rodenburg, 2011). Final OCRs were calculated as follows: non-mitochondrial OCR was subtracted from Routine, Leak and ETC values; the resulting values were normalized to CS activity per gram of protein.

### SDS-PAGE and Western blotting

Protein samples were separated on 4-20% Criterion TGX Stain-Free protein gels (Bio-Rad; 5671093) or regular 10% acrylamide gels using Laemmly SDS-PAGE. The separated proteins were electrophoretically transferred to nitrocellulose membranes (Bio-Rad) and incubated with the corresponding primary and HRP-conjugated secondary antibodies for protein detection. Western blots were detected with ChemiDoc XRS+ Imaging System (Bio-Rad).

### Statistical analysis and data visualization

For the IF particle analysis, relative mtDNA copy number quantification and high-resolution respirometry, an unpaired two-tailed Student’s t-test was used to test the null hypothesis of no changes between the conditions. Statistical significance was determined using a threshold of *p* ≤ 0.05. The statistical analysis for BioID was different and is described in the respective section.

Bar charts and volcano plots were created in R-studio, the Venn diagram was created in BioVenn software (Hulsen et al., 2008), heatmaps for complexome profiling were created in Microsoft Excel, and microscopic images were processed with ImageJ and Adobe Illustrator.

## Data availability

Proteomics raw files and MaxQuant output files for the BioID experiment are deposited in ProteomeXchange (www.proteomexchange.org, PRIDE accession number PXD037857). Results of complexome profiling are deposited in CEDAR database (van Strien et al., 2021), a resource dedicated for storage and sharing of complexomics datasets (www3.cmbi.umcn.nl, accession number CRX38). The presented data can be found in the paper or supplementary data. Additional data is available upon request from the corresponding authors.

## Author contributions

AP, MH and JNS initially conceived and designed this study with subsequent contributions in the experimental design by SG and AC-O. Experimental work was carried out by AP, AC-O and AH. AC-O and AP performed LC/MS-MS analysis. All authors were involved in data analysis and interpretation of the results. The first draft of the manuscript was written by AP with all authors contributing to revisions and the final draft of the manuscript, which has been approved by all authors.

## Acknowledgements

We thank Prof. Ulrich Brandt for access to native electrophoresis and MS, Helga van Rennes for laboratory assistance, Dr. Daan Pannemaan and Dr. Richard Rodenburg for assistance in knockout generation, the Spiegroup of TML for enzyme activity measurements, Rafail Kyranas for help with immunofluorescence experiments, and Femke Vermeir for help with cloning and BioID.

AP and JNS were supported by the European Union’s Horizon 2020 research and innovation program under the Marie Sklodowska-Curie grant agreement No 721757. AC-O and MH were supported by the Netherlands Organization for Health Research and Development (ZonMW TOP 91217009). A.H. and S.G. were supported by the Academy of Finland (grant No 332458).

