## Supplementary Figures for "Uncharacterized protein c17orf80: a novel interactor of human mitochondrial nucleoids"

**A**

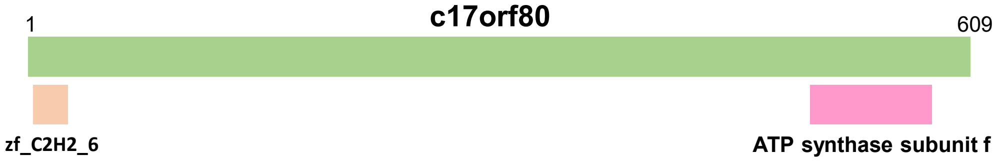


```
6J5I_f ATP synthase subunit f, mitochondrial; membrane protein; HET: ADP, ATP; 3.34A {Sus scrofa}
Probability: 99.54%,   E-value: 2.6e-15,   Score: 128.93,   Aligned cols: 82,   Identities: 26%,   Similarity: 0.437,   Template Neff: 9.3
Q ss_pred                  ccCCccccccccccCCCCchhhcCCCCchHHHHHHHHHheeE---EEeeecCCCccHHHHHHhheeEeeeeeccchHHHH-hhcCCC
Q ENSP0000044055    503  SQVSLLERSSTHIRSLEPPAGLTTSNFSLMRLLGAVQKGWIR---CNTTIRKSGFGGITMLFTGYFVLCCSWSFRRLKK-LCRPLP   584 (609)
Q Consensus        503  ~qgpL~eRsl~dVRl~ElP~WLat~~~S~pRll~avqk~W~w---YYidVKKGGVGGItMLlAGYCVLsY~WSY~HlK~-~WRkyh   584 (609)
                          +++|+.++.++||+++|+|.||..++++..+...++.+.|.+ -||++++++++||.|+++|+|||+|+|+|+|++. +|+|||
T Consensus        2    ~~~~~~~~~~~~~k~~e~p~w~~~r~~~~~~~~~~~~r~~~~y~~ky~~~~~~~~~~~~~~~~~~~y~~~~y~~~~~~~~~~~~~kyh   87 (87)
T 6J5I_f           2    SVVPLKDRRLLEVKLGELPSWILMRDFTPSGIAGAFQRGYYRYYNKYVNVKKGSVAGLSMVLAAYVVFNYCRSYKELKHERLRKYH   87 (87)
T ss_dssp                   CHHHHHHHHHHHHHHHTCC-------CCTTHHHHCSHHHHGGGSCSCCCCTTTSCHHHHTTHHHHHHHHHHHHHHTTTCCCCCSSCCC
T ss_pred                   CCCCCCCCCccccCccccchhHHHhccCCCHHHHHHHHHHhchhhhhhcCcCCCChHHHHHHHHHHHHHHHHHHhcHHHHHHHhccC
```

```
PF18450.4 zf_C2H2_6 ; Zinc Finger domain
Probability: 91.33%,   E-value: 0.097,   Score: 37.16,   Aligned cols: 21,   Identities: 33%,   Similarity: 0.729,   Template Neff: 4.1
Q ss_pred                         ccCCCCCCchhh---cccCCCcccc
Q ENSP0000044055           9      EVCPYCKKPFKR---LKSHLPYCK     29 (609)
Q Consensus                9      EvCPyCgKpFKR---LKSHLPHCK     29 (609)
                                 .+||||++.|-+ ++-|.-||.
T Consensus                3      ~~CpyC~lrF~sp~~k~~He~~C~     26 (28)
T PF18450.4                3      TVCIYCSLRFSSLELKHDHEITCE     26 (28)
T ss_pred                          ccccccCCccccChHHHHHHhhhcc
```

**B**

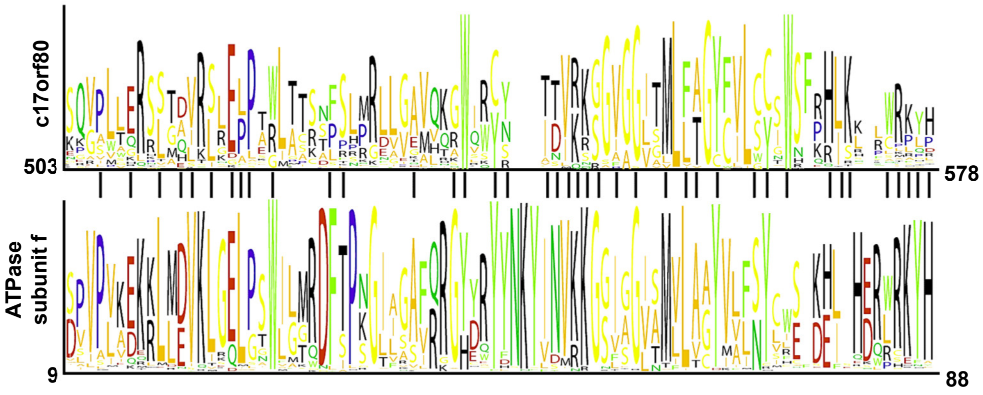


**Figure S1. c17orf80 homology analysis. A.** HHPred search results. Schematic representation of detected homology domains followed by alignments of c17orf80 with ATP synthase subunit f and a zinc finger domain. **B**. Sequence logos of c17orf80 C-terminus and vertebrate ATP synthase subunits f showing conservation of the region. Vertical lines between the template and query indicate similar and identical residues.

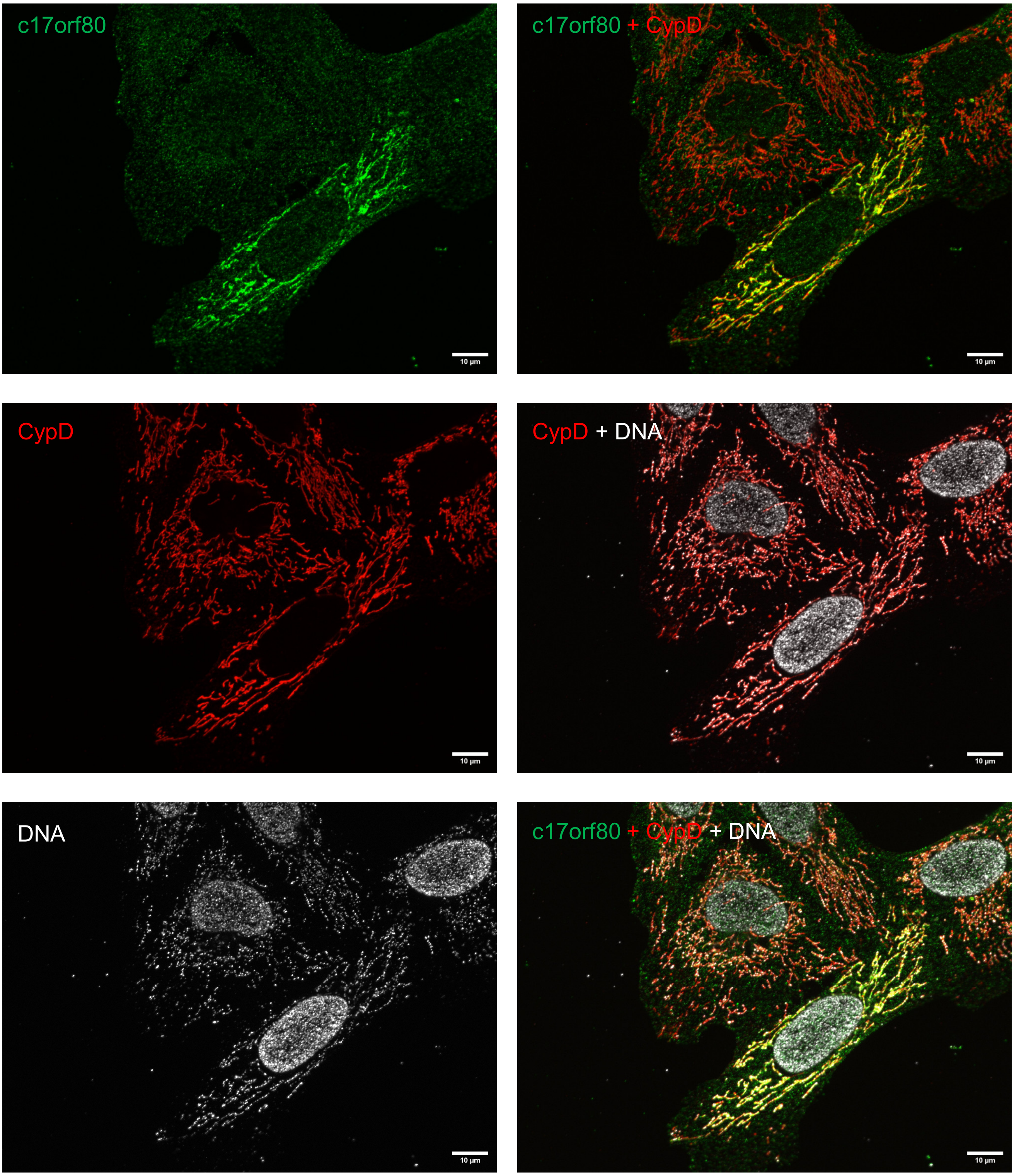


**Figure S2. siRNA-mediated silencing of c17orf80.** Cells with successful (top) and unsuccessful (bottom) knockdowns for c17orf80 are shown for comparison of IF signals. C17orf80 immunofluorescence was observed in the mitochondrial network and cytoplasm, however, the cytoplasmic signal does not relate to c17orf80. Co-immunostaining with c17orf80, CypD (mitochondrial network), and DNA (nucleoids and nuclei) antibodies in U2OS cells. The scale bar is 10 µm.

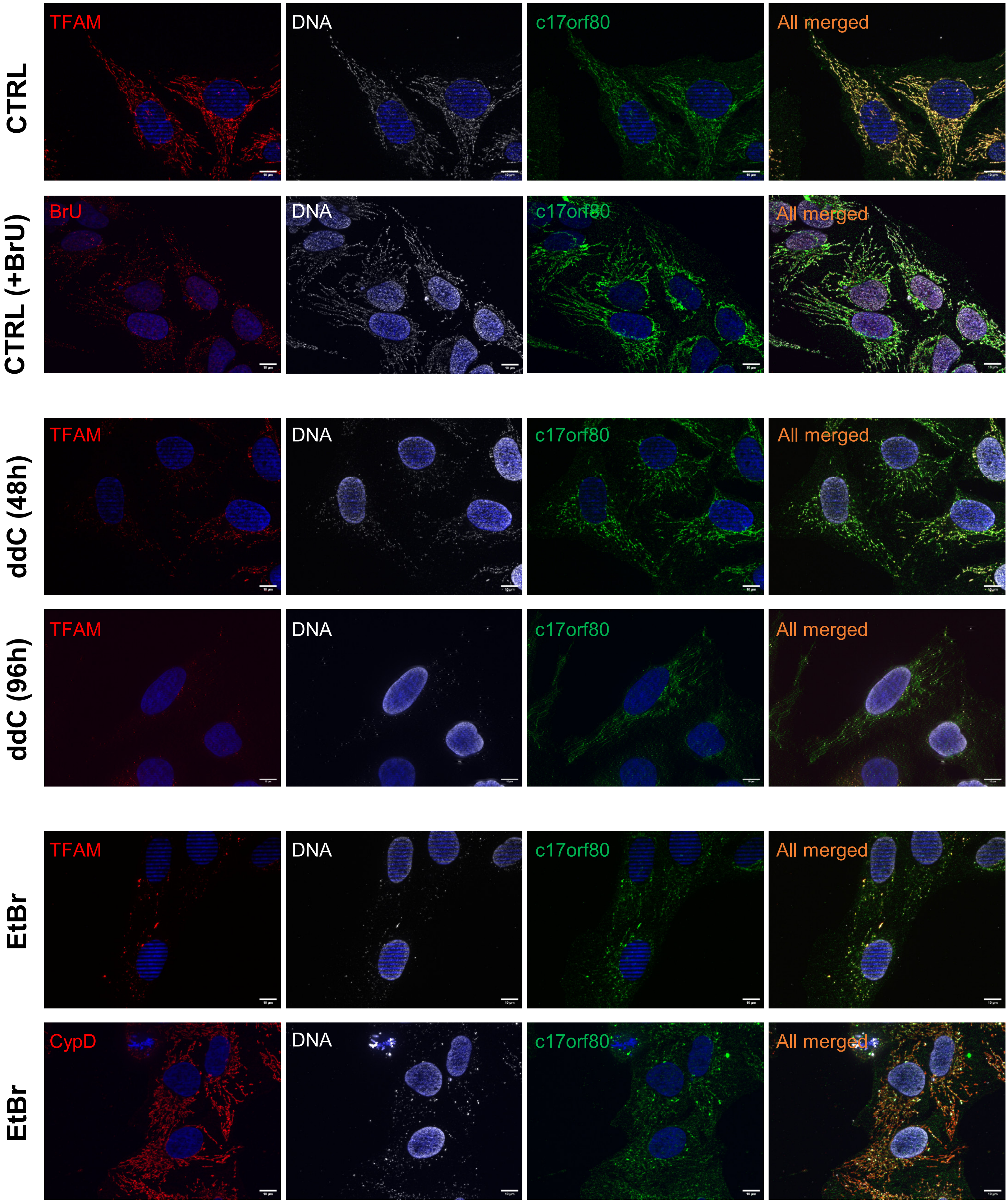


**Figure S3. C17orf80 co-localizes with mtDNA in U2OS cells.** Uncropped images for the main Figure 4. Co-immunostaining for c17orf80, TFAM (nucleoids), CypD (mitochondrial network), DNA (nucleoids and nuclei) or BrU (RNA-granules) antibodies in U2OS cells**.** Nuclei were stained with DAPI. The scale bar is 10 µm.

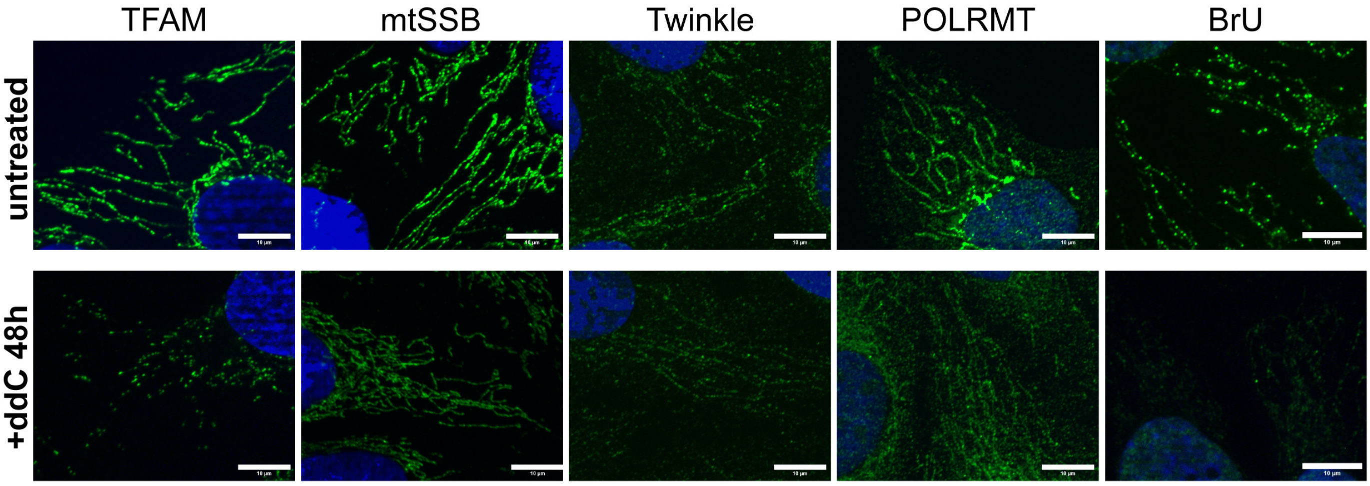


**Figure S4. Nucleoid-associated proteins and RNA granules before and after ddC treatment.** IF images of ddC-treated and control cells stained with antibodies against TFAM, mtSSB, Twinkle or POLRMT, or anti-BrU (after 1 h BrU-labelling). Nuclei stained with DAPI (blue). The scale bar is 10 µm.

**c17orf80-BirA-C**

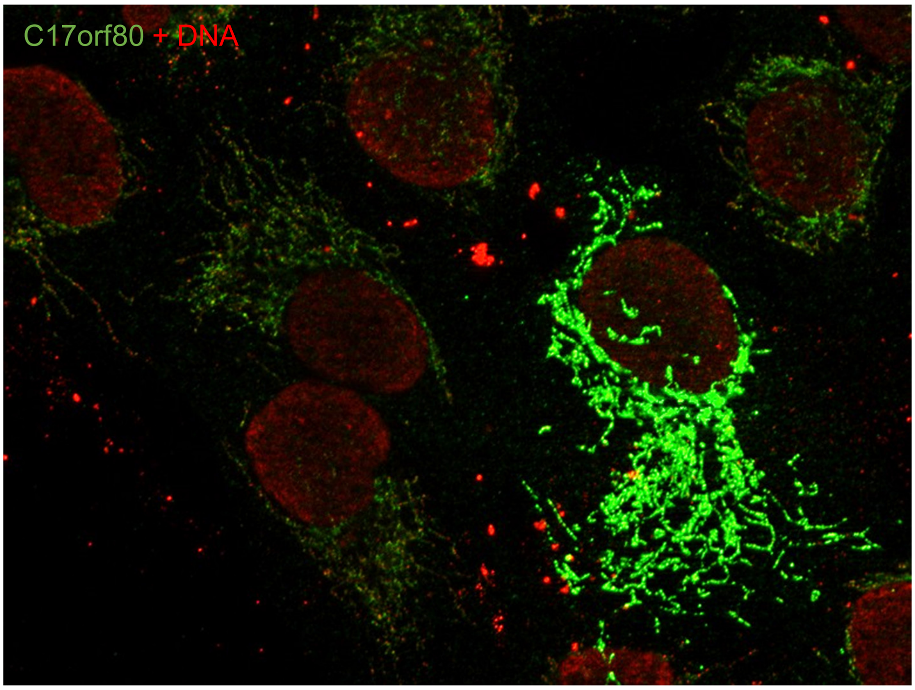


**c17orf80-BirA-N**

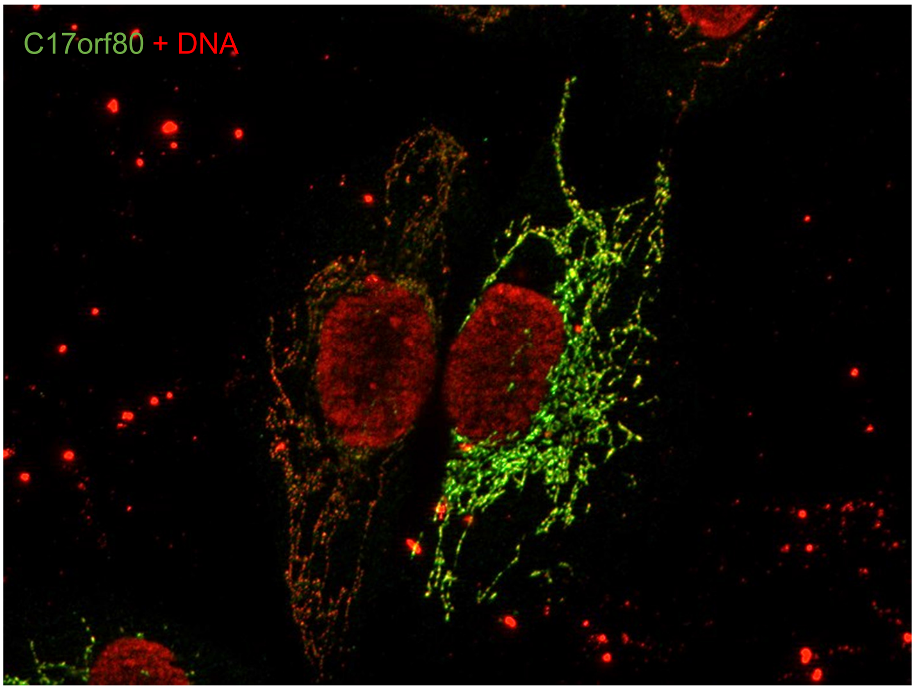


**Figure S5. Both C- and N-terminally tagged c17orf80-BirA* fusion proteins were imported to mitochondria.** Co-immunostaining for c17orf80 and DNA (nucleoids and nuclei) in U2OS cells overexpressing the fusion protein after transient transfection with pDEST-pcDNA5-c17orf80-BirA*-FLAG N- or C-term plasmids.

**A**

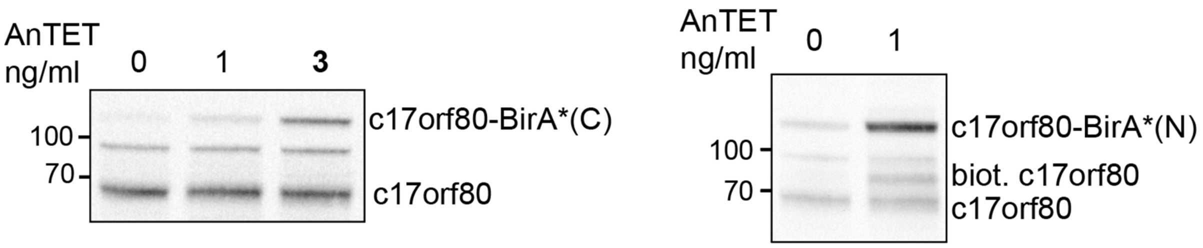


**B**

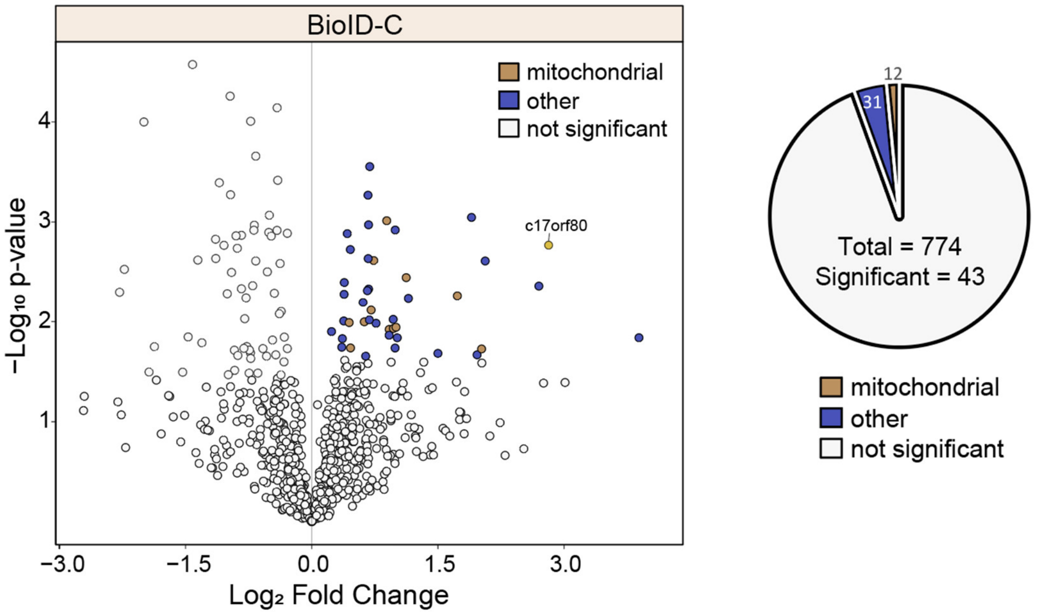


**Figure S6. Results of c17orf80 interactome mapping with c17orf80-BirA*(C) fusion. A.** Levels of c17orf80-BirA* fusion proteins after induction with AnTET. The Western blot was detected with the anti-c17orf80 antibody. **B.** Volcano plot depicts average log2-transformed intensity fold changes plotted against negative log10-transformed P values for four biological replicates. Only 43 proteins were significantly enriched in c17orf80-BirA*(C) pulldown out of which 12 were mitochondrial. Coloured points indicate proteins that were significantly enriched in the c17orf80-BirA*(C) pulldown compared to the non-induced control. Fold change was defined as a ratio of c17orf80-BirA*(N) to control. Statistical significance was determined by FDR < 0.05 with S0 = 0.1.

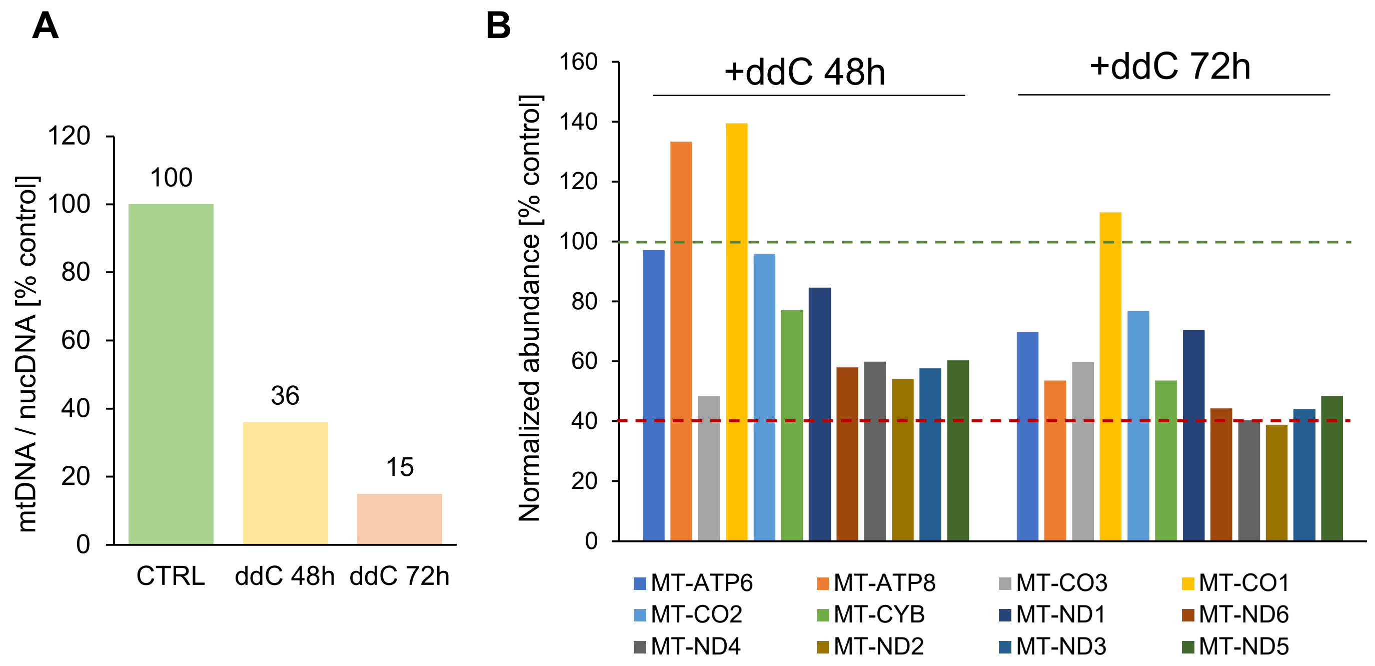


**Figure S7. Effect of ddC treatment on mtDNA content and mtDNA-encoded OXPHOS subunits abundance in the CP samples. A.** The relative mtDNA copy number decreased to 36% and 15% of that in control after 48 h and 72 h of treatment with 100 µM ddC as measured by qPCR (n = 1). **B**. Levels of the mtDNA-encoded OXPHOS subunits after ddC-treatment detected in the CP dataset. Total iBAQ intensities of ddC-treated samples are shown in relation to that of the control. MT-ND4L was not identified by MS in this dataset.

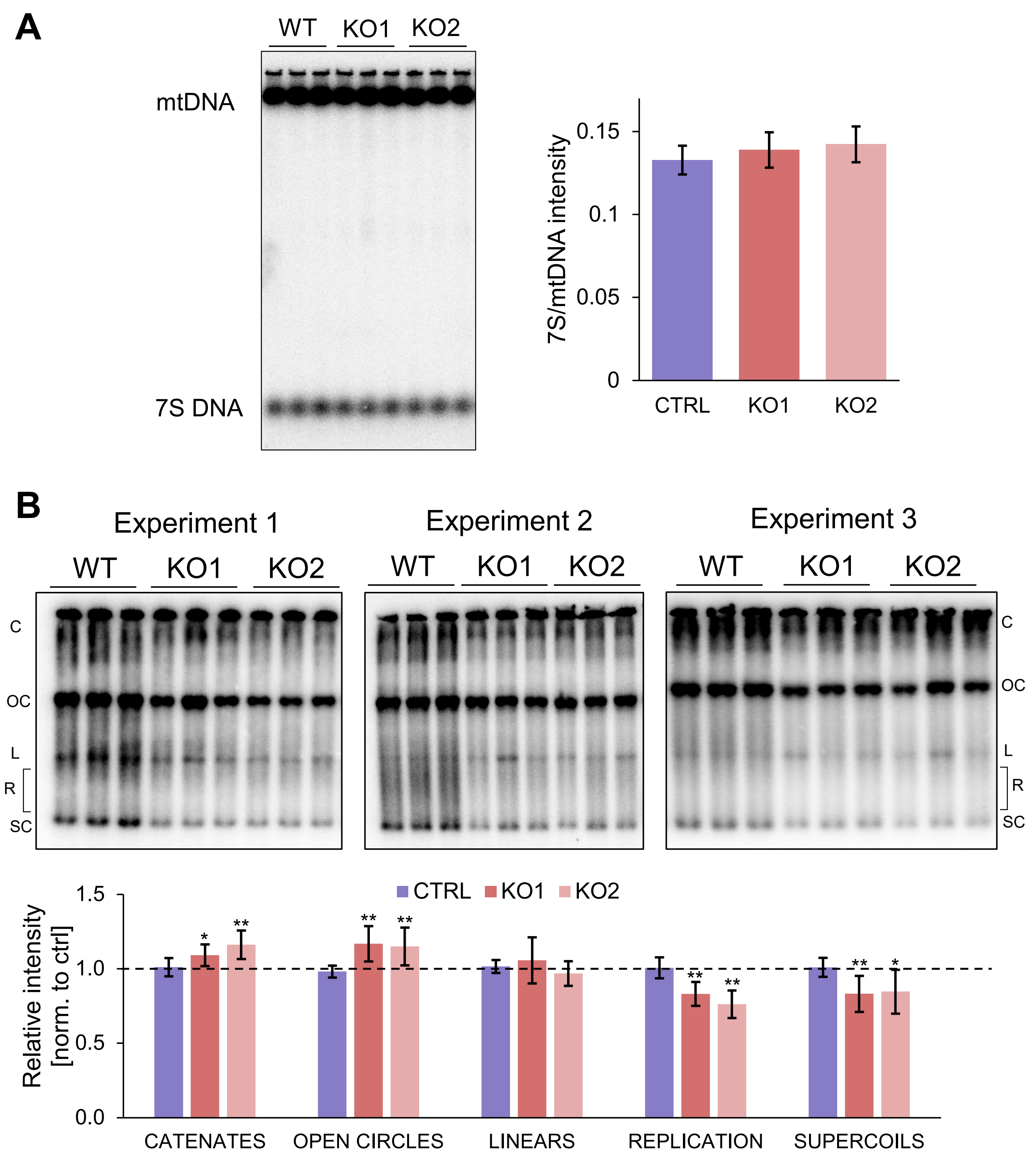


**Figure S8. A.** Analysis of topological forms of mtDNA. KOs showed a 10-15% increase in catenates and open circles and a 15-25% decrease in supercoiled and replicating mtDNA molecules. Three independent Southern blots each containing three biological replicates of each cell line were analysed. The intensity of each topological form was measured in relation to the total intensity of the line. The first replicate of control was set to 1 for every topological form in each experiment. Data are mean ± SD of fold changes between control and KO from nine biological replicates; unpaired Student's t-test. Cutoffs for statistical significance: *P ≤ 0.05, **P ≤ 0.01. C = catenates (oligomeric mtDNA), OC = open circles, L = linear, R = replication, SC = supercoils. **B.** 7S DNA Southern blot and its quantification. No significant changes in 7S DNA levels were observed. Data are mean ± SD of the ratio between 7S DNA and mtDNA intensities of three biological replicates; unpaired Student's t-test.

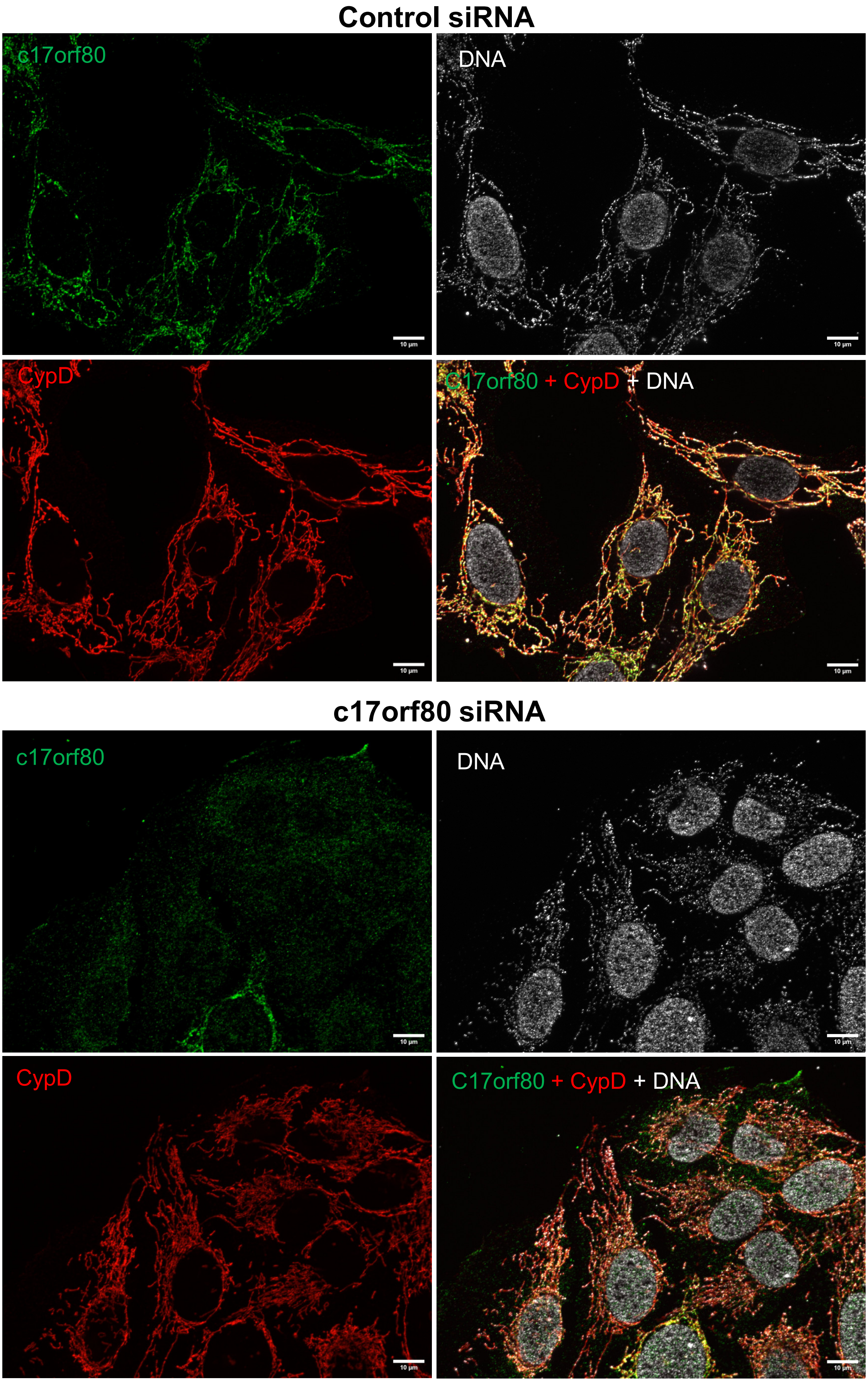


**Figure S9. The morphology of mitochondrial network and nucleoids does not change after depletion of c17orf80.** Cells were treated with c17orf80-targeted siRNA or a control siRNA for three days. Co-immunostaining with c17orf80, CypD (mitochondrial network), and DNA (nucleoids and nuclei) antibodies in U2OS cells. The scale bar is 10 μm.
