## Supplementary figures and images for "Uncharacterized protein c17orf80: a novel interactor of human mitochondrial nucleoids"

### Supplementary Data 1

*Supplementary Data 1:*
*Immunofluorescene images of control and ddC-treated U2OS cells.*

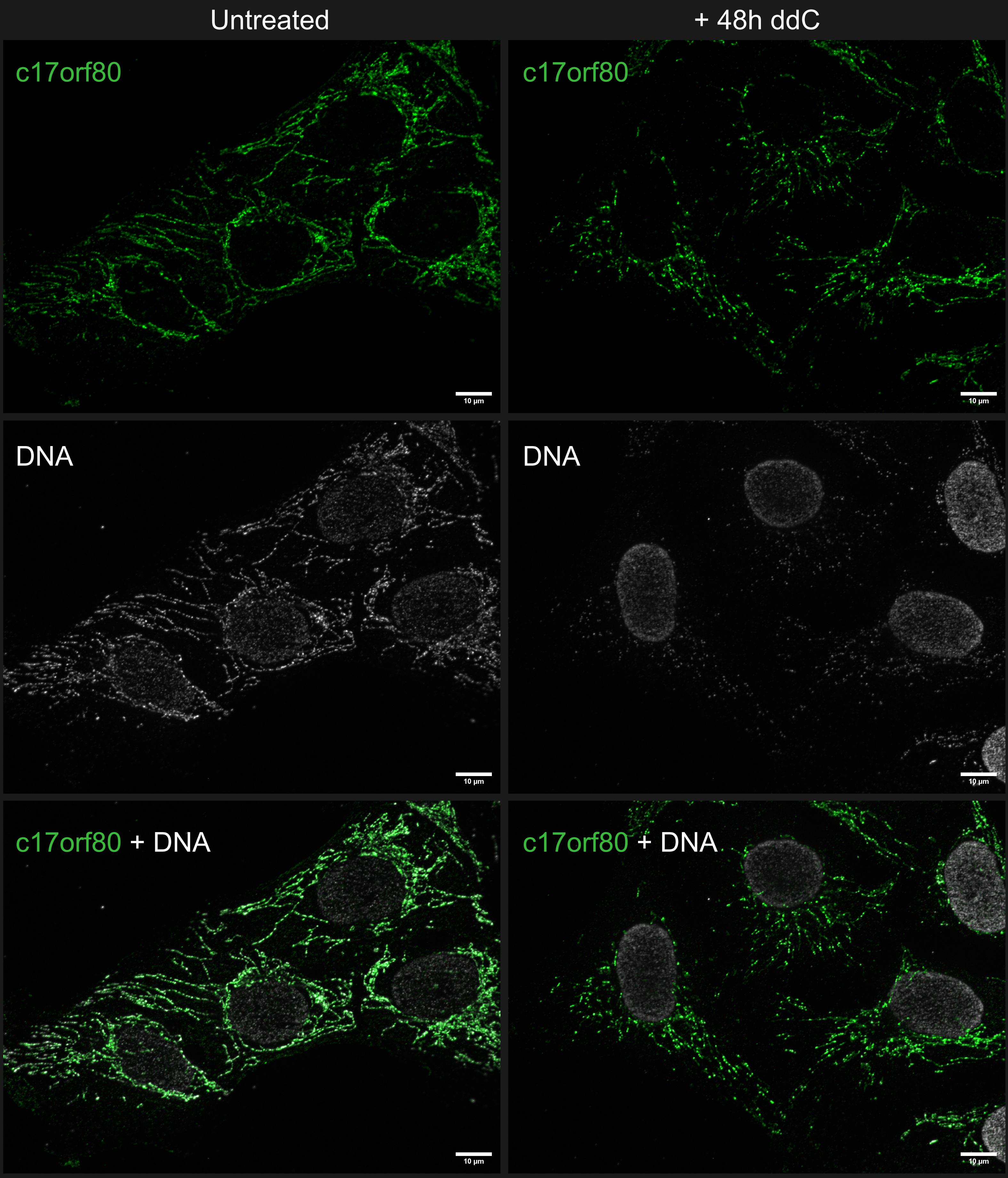

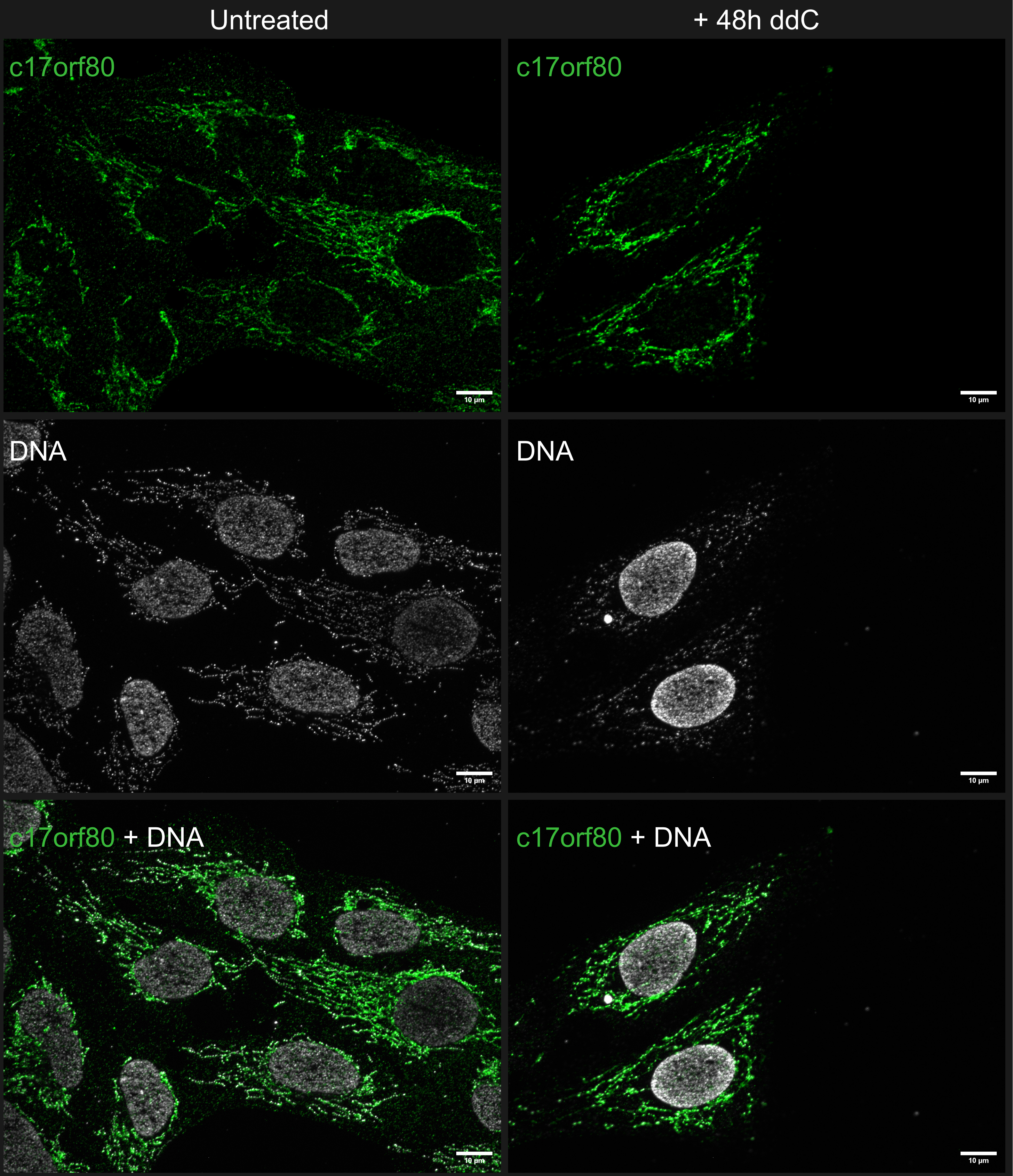

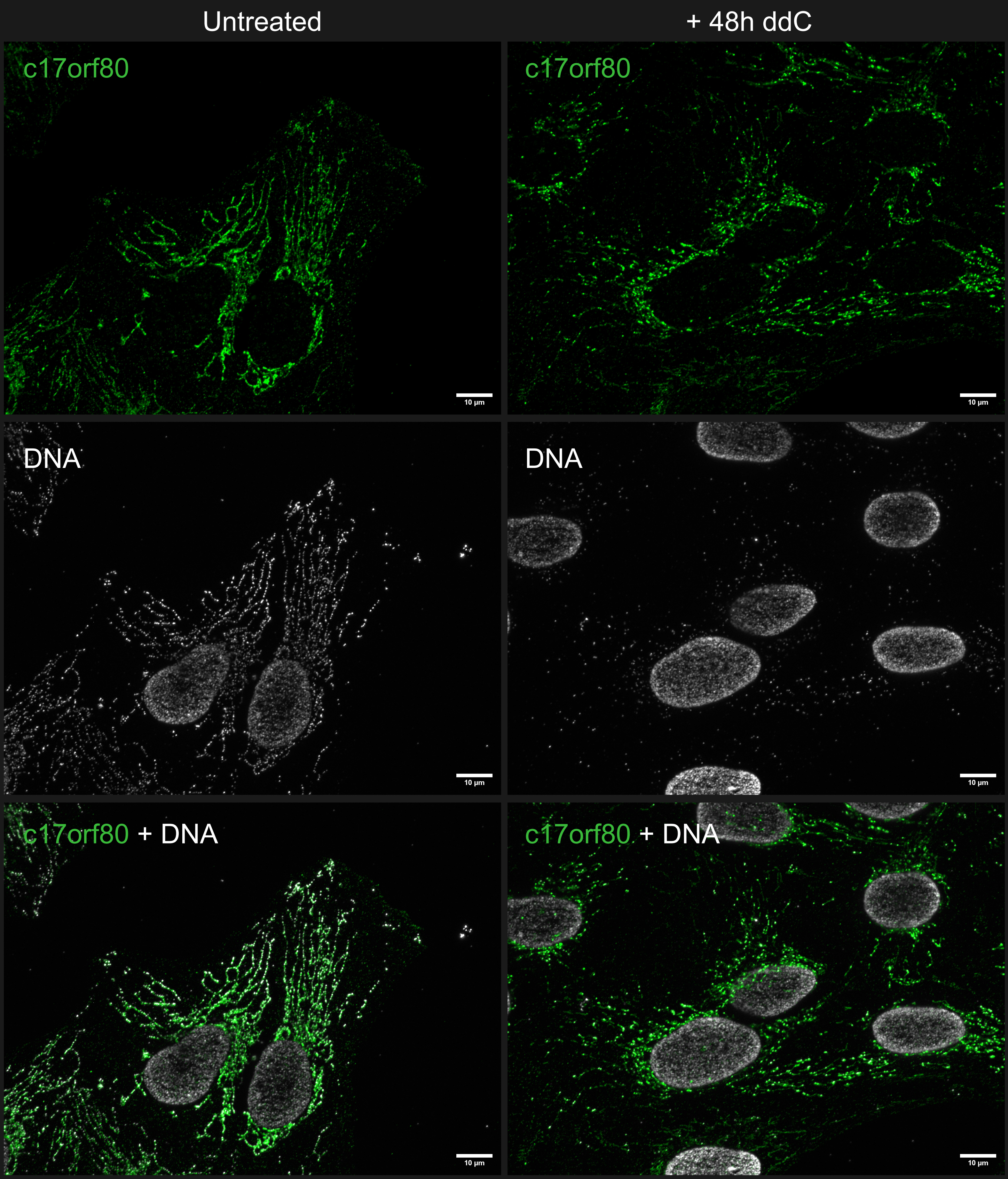

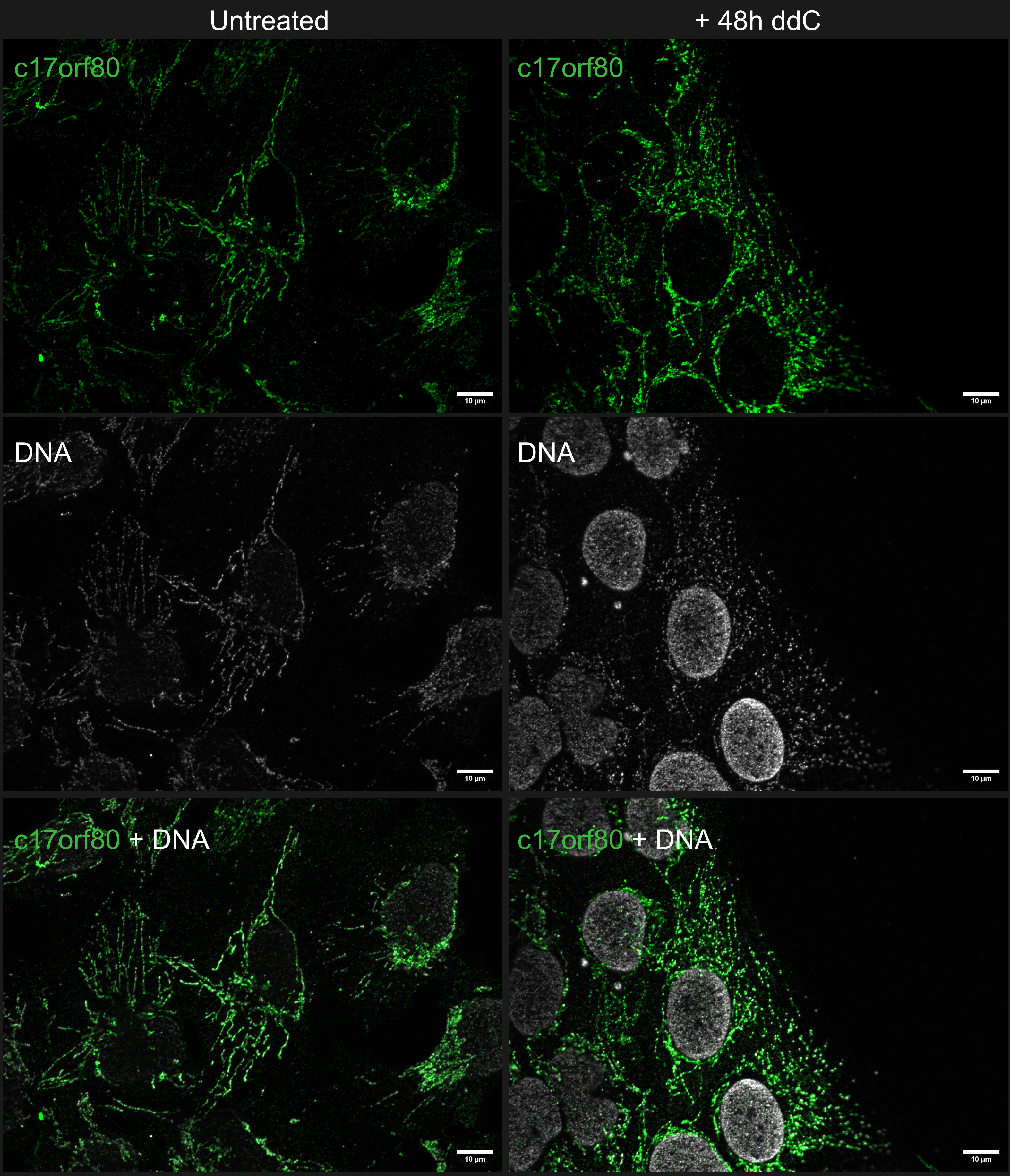

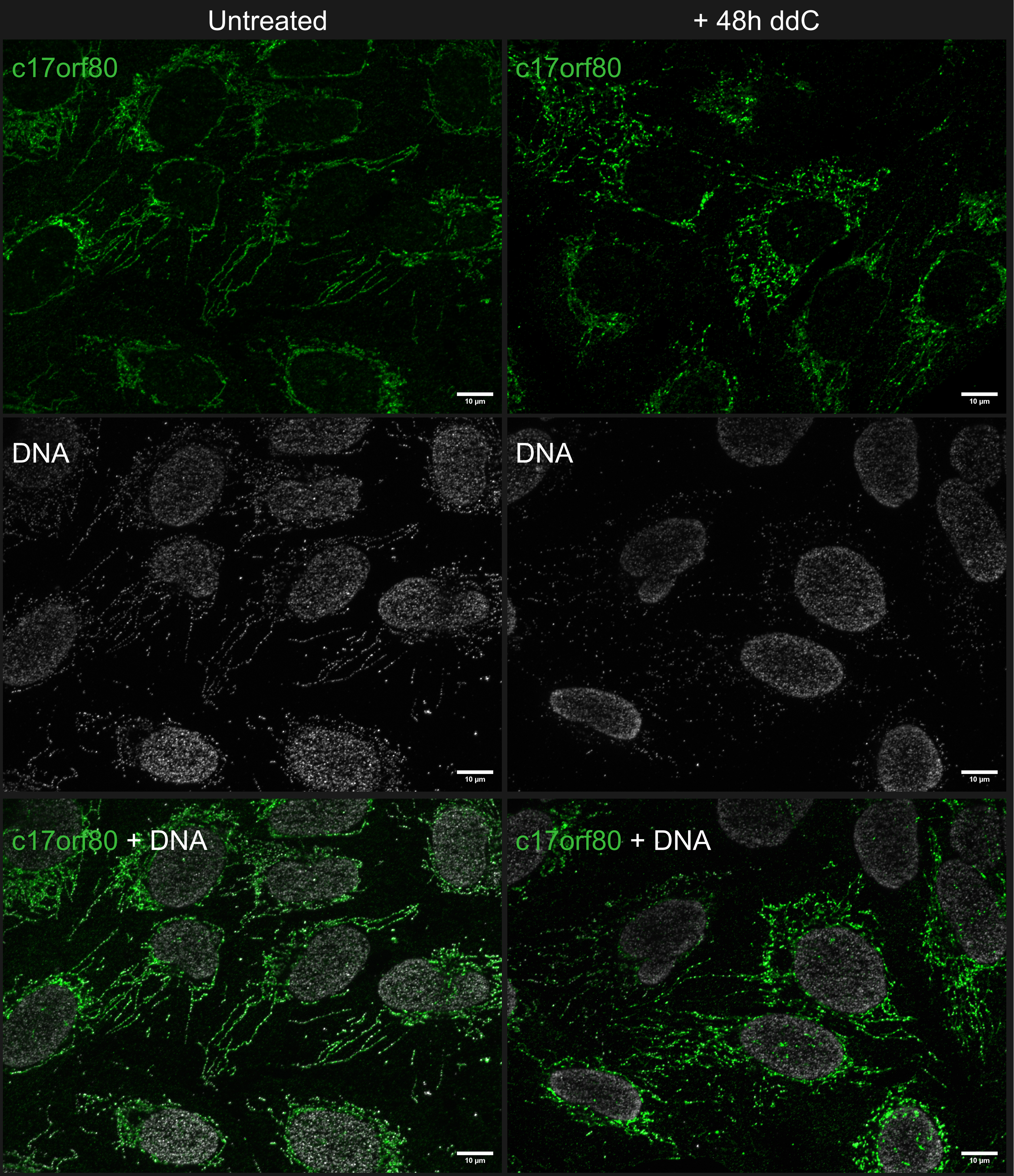
